# IFNγ regulates MR1 transcription and antigen presentation

**DOI:** 10.1101/2025.02.09.637183

**Authors:** ME Huber, EA Larson, TN Lust, CM Heisler, MJ Harriff

**Author notes:** Author contributions: MEH, EAL, TNL, and CMH performed experiments. MEH and MJH designed the experiments and wrote the manuscript. MEH conducted data analysis and prepared the figures. All authors reviewed the manuscript.

## Abstract

Antigen presentation molecules play key roles in activating T cell immunity. Multiple complementary pathways are known to regulate classical MHC-I molecules at transcriptional, translational, and post-translational levels. Intracellular trafficking mechanisms dictating post-transcriptional regulation of MR1, the MHC Class I-like molecule which restricts MAIT cells, have been an area of focus; however, little is known about *MR1* transcriptional regulation. We demonstrate that, similar to classical MHC-I, interferons regulate *MR1* transcription. Treatment of airway epithelial cells (AEC) with recombinant IFNβ or IFNγ variably increased *MR1* transcripts, while only IFNγ significantly increased surface MR1 expression and enhanced antigen presentation to MAIT cells. The MR1 promoter contains binding motifs for interferon regulatory factor 1 (IRF1), an important MHC-I transcription factor. IRF1 knockout reduced IFNγ-stimulated MR1 transcription, surface expression, and antigen presentation. Conversely, knockout of Nod-like Receptor family CARD domain containing 5 (NLRC5), a critical component of IFNγ-induced MHC-I transcription, did not significantly impact MR1 expression. These findings were corroborated in primary human AEC treated with IFNγ. In co-culture experiments, MAIT cells incubated with *Streptococcus pneumoniae*-infected primary AEC produced sufficient IFNγ to stimulate upregulation of MR1 expression. Our data support a model where IFNγ from activated MAIT cells or another source stimulates IRF1-dependent MR1 expression and antigen presentation, leading to greater MAIT cell activation. A robust MR1-dependent MAIT cell response may be beneficial for early infection responses, allowing minimal antigen stimulus to generate greater proinflammatory activity.

## Introduction

Mucosal-associated invariant T (MAIT) cells, an innate-like subset of T lymphocytes that comprise a relatively large proportion of the total CD8^+^ T cell population in human blood and lungs, play key roles in clearing respiratory bacterial, fungal, and viral infections^1–3^. Upon antigen presentation, MAIT cells are capable of immediate effector function and release inflammatory cytokines like interferon-γ (IFNγ) and tumor necrosis factor (TNFα)^2–5^. This rapid activation primes MAIT cells to coordinate early infection response, but also necessitates tight regulation of antigen presentation to prevent inappropriate MAIT cell activation to inappropriate stimuli.

MAIT cells are restricted by the MHC class I-related molecule MR1, which presents small molecule metabolite antigens such as those generated during bacterial riboflavin biosynthesis^2,3,6,7^. There is a large pool of potential MR1 ligands produced by commensal airway flora in addition to pathogenic respiratory microbes. *MR1* mRNA is expressed across cell types and tissues, and MR1 proteins primarily reside in intracellular compartments like the ER and endosomal compartments^7–11^. The basal intracellular location of MR1 and ligand-induced translocation to the cell surface play critical parts in regulation of MAIT activation (as reviewed in ^12–14^).

The intracellular trafficking mechanisms dictating post-transcriptional regulation of MR1 have been an area of research focus; however, little is known about *MR1* transcriptional regulation. Multiple complementary pathways regulate classical MHC-Ia molecules at transcriptional, translational, and post-translational levels^15,16^. Interferons (IFNs) like IFNβ and IFNγ drive transcription of MHC-Ia through expression of downstream transcription factors like Interferon Regulatory Factor 1 (IRF1) and Nod-like receptor family CARD domain containing 5 (NLRC5), which in turn bind to the *HLA* promoter to induce transcription^15–19^. Although the *MR1* gene resides on human chromosome 1, outside the chromosome 6 *HLA* locus^7^, these pathways may provide insight into transcriptional regulation of MR1. Recent research links *MR1* expression with disease pathology (e.g. meningeal tuberculosis^20^, glioma^21^, and COPD^22–24^), although specific mechanisms controlling *MR1* transcription remain unclear.

Here, we investigated the role of IFNγ in stimulating MR1 expression in human airway epithelial cells (AEC). We found IFNγ promotes MR1 transcription, antigen presentation, and MAIT cell responses. While NLRC5 and IRF1 were both important for IFNγ-induced *HLA-A* transcription, NLRC5 was largely dispensable for *MR1* transcription. Finally, we demonstrate that MAIT cells, activated in co-culture with infected AEC, produce sufficient IFNγ to stimulate *MR1* transcription. Taken together, our data support a model in which IFNγ from activated immune cells induces MR1 expression and antigen presentation, leading to greater MAIT cell activation. These results establish an additional level of MR1 regulation, informing our understanding of MAIT cell activation and dysregulation in infection and disease.

## Results

### MR1 expression increases in infected AECs co-cultured with MAIT cells

Previously, our lab examined the expression of MR1 in primary human AEC from donors with chronic obstructive pulmonary disease (COPD) or history of tobacco use^22^. MAIT cell responses to these primary AEC infected with *Streptococcus pneumoniae* (*Sp*) were reduced in the contexts of COPD and smoking. These altered MAIT cell responses were MR1-dependent, but we observed minimal differences in the baseline *MR1* mRNA expression of these primary AEC alone. We therefore hypothesized that co-culture with MAIT cells could impact MR1 expression and function. To address this, we examined *MR1* mRNA expression of primary AEC co-cultured with MAIT cells alone or in the context of *Sp* infection.

We noticed significantly increased *MR1* mRNA expression in AEC from healthy donors when infected with *Sp* and cultured with MAIT cells (Figure 1A, Supplemental Figure 1A, Table 1). Infection with *Sp* or co-culture with MAIT cells alone did not stimulate a significant response. We replicated this system using a model bronchial epithelial cell line (BEAS-2B cells) infected with *Mycobacterium smegmatis* (*Ms*). Similarly, we observed increased *MR1* expression in the *Ms*-infected BEAS-2B cells co-cultured with MAIT cells, with no impact of either condition alone (Figure 1B, Supplemental Figure 1B, Table 1). Using flow cytometry to quantify surface MR1 protein expression, we likewise found increased MR1 expression with both *Ms* infection and MAIT cell co-culture compared to either condition alone (Figure 1C, Supplemental Figure 1C). These data suggest that MAIT cells, when activated by presentation of bacterial antigens, could lead to increased *MR1* mRNA or surface protein expression in the infected cell.

**Figure 1.**
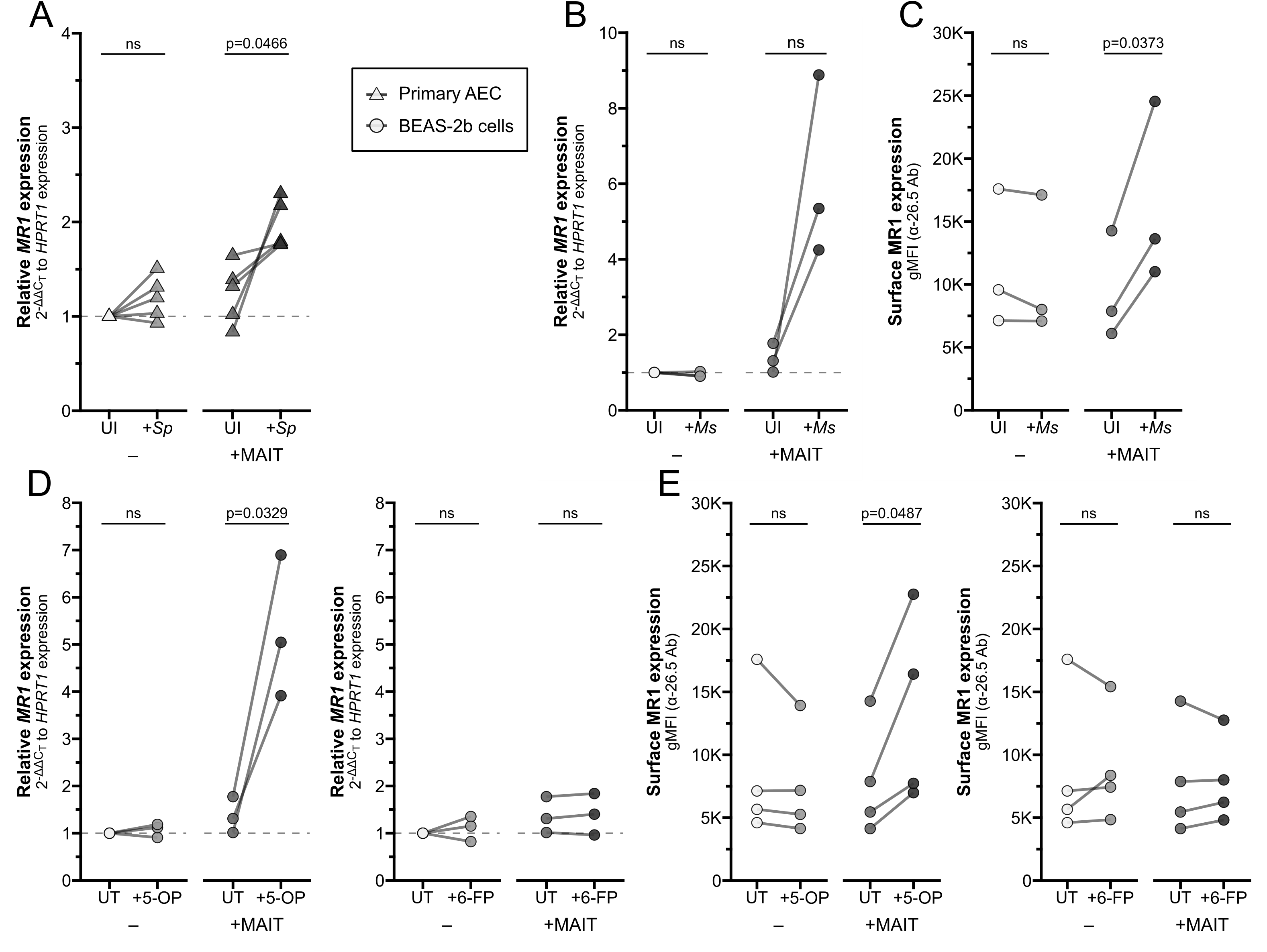
Increased MR1 expression following MAIT cell activation. **(A)** RT-qPCR of RNA isolated from primary human AECs (n=5) infected with *S. pneumoniae* (*Sp*) for one hour and incubated overnight with MAIT cell clone. *MR1* expression was calculated relative to *HPRT1* expression and uninfected no-MAIT (UI-) controls, paired by individual donor. MR1 **(B)** mRNA and **(C)** surface expression of BEAS-2B cells infected with *M. smegmatis* (*Ms*) for one hour and incubated overnight with MAIT cell clone. **(B)** RT-qPCR of *MR1* expression was calculated relative to *HPRT1* expression and UI-control, paired by experimental replicate. **(C)** Geometric mean fluorescence intensity (gMFI) of surface MR1 stained with α-MR1 26.5 Ab, paired by experimental replicate. MR1 **(D)** mRNA and **(E)** surface expression of BEAS-2B cells treated with 5-OP-RU (left, “5-OP”) or 6-FP (right) for one hour and incubated overnight with MAIT cell clone. **(D)** RT-qPCR of *MR1* expression was calculated relative to *HPRT1* expression and UT-control, paired by experimental replicate. **(E)** gMFI of surface MR1 stained with α-26.5 Ab, paired by experimental replicate. Pairwise statistical analyses are in Table 1.

**Table 1.** Statistics associated with Figure 1 and Supplemental Figure 1.

| Figure | Data | Sample 1 |  |  | Sample 2 |  |  | n1 | n2 | df | statistic | p-value | sig. |
| --- | --- | --- | --- | --- | --- | --- | --- | --- | --- | --- | --- | --- | --- |
|  |  | Cell | Ag | T cell | Cell | Ag | T cell |  |  |  |  |  |  |
| 1A, SF1A | *MR1* mRNA | AEC | UI | NT | AEC | *Sp* | NT | 5 | 5 | 4 | 1.913 | 0.1283 | ns |
| 1A, SF1A | *MR1* mRNA | AEC | UI | NT | AEC | UI | MAIT | 5 | 5 | 4 | 1.702 | 0.1639 | ns |
| 1A, SF1A | *MR1* mRNA | AEC | UI | NT | AEC | *Sp* | MAIT | 5 | 5 | 4 | 8.332 | 0.0011 | ** |
| 1A, SF1A | *MR1* mRNA | AEC | *Sp* | NT | AEC | UI | MAIT | 5 | 5 | 4 | 0.391 | 0.7157 | ns |
| 1A, SF1A | *MR1* mRNA | AEC | *Sp* | NT | AEC | *Sp* | MAIT | 5 | 5 | 4 | 3.679 | 0.0212 | * |
| 1A, SF1A | *MR1* mRNA | AEC | UI | MAIT | AEC | *Sp* | MAIT | 5 | 5 | 4 | 2.846 | 0.0466 | * |
| 1B, SF1B | *MR1* mRNA | B2B | UI | NT | B2B | *Ms* | NT | 3 | 3 | 2 | 1.387 | 0.2998 | ns |
| 1B, SF1B | *MR1* mRNA | B2B | UI | NT | B2B | UI | MAIT | 3 | 3 | 2 | 1.653 | 0.24 | ns |
| 1B, SF1B | *MR1* mRNA | B2B | UI | NT | B2B | *Ms* | MAIT | 3 | 3 | 2 | 3.689 | 0.0663 | ns |
| 1B, SF1B | *MR1* mRNA | B2B | *Ms* | NT | B2B | UI | MAIT | 3 | 3 | 2 | 2.253 | 0.153 | ns |
| 1B, SF1B | *MR1* mRNA | B2B | *Ms* | NT | B2B | *Ms* | MAIT | 3 | 3 | 2 | 3.702 | 0.0658 | ns |
| 1B, SF1B | *MR1* mRNA | B2B | UI | MAIT | B2B | *Ms* | MAIT | 3 | 3 | 2 | 3.095 | 0.0905 | ns |
| 1C, SF1C | αMR1 gMFI | B2B | UI | NT | B2B | *Ms* | NT | 3 | 3 | 2 | 1.31 | 0.3205 | ns |
| 1C, SF1C | αMR1 gMFI | B2B | UI | NT | B2B | UI | MAIT | 3 | 3 | 2 | 0.8576 | 0.4815 | ns |
| 1C, SF1C | αMR1 gMFI | B2B | UI | NT | B2B | *Ms* | MAIT | 3 | 3 | 2 | 15.02 | 0.0044 | ** |
| 1C, SF1C | αMR1 gMFI | B2B | *Ms* | NT | B2B | UI | MAIT | 3 | 3 | 2 | 0.7913 | 0.5117 | ns |
| 1C, SF1C | αMR1 gMFI | B2B | *Ms* | NT | B2B | *Ms* | MAIT | 3 | 3 | 2 | 12.17 | 0.0067 | ** |
| 1C, SF1C | αMR1 gMFI | B2B | UI | MAIT | B2B | *Ms* | MAIT | 3 | 3 | 2 | 5.032 | 0.0373 | * |
| 1D, SF1D | *MR1* mRNA | B2B | UT | NT | B2B | 5-OP | NT | 3 | 3 | 2 | 0.8515 | 0.4842 | ns |
| 1D, SF1D | *MR1* mRNA | B2B | UT | NT | B2B | UT | MAIT | 3 | 3 | 2 | 1.653 | 0.24 | ns |
| 1D, SF1D | *MR1* mRNA | B2B | UT | NT | B2B | 5-OP | MAIT | 3 | 3 | 2 | 4.931 | 0.0388 | * |
| 1D, SF1D | *MR1* mRNA | B2B | 5-OP | NT | B2B | UT | MAIT | 3 | 3 | 2 | 1.635 | 0.2437 | ns |
| 1D, SF1D | *MR1* mRNA | B2B | 5-OP | NT | B2B | 5-OP | MAIT | 3 | 3 | 2 | 4.784 | 0.041 | * |
| 1D, SF1D | *MR1* mRNA | B2B | UT | MAIT | B2B | 5-OP | MAIT | 3 | 3 | 2 | 5.377 | 0.0329 | * |
| 1D, SF1D | *MR1* mRNA | B2B | UT | NT | B2B | 6-FP | NT | 3 | 3 | 2 | 0.7058 | 0.5535 | ns |
| 1D, SF1D | *MR1* mRNA | B2B | UT | NT | B2B | 6-FP | MAIT | 3 | 3 | 2 | 1.585 | 0.2539 | ns |
| 1D, SF1D | *MR1* mRNA | B2B | 6-FP | NT | B2B | UT | MAIT | 3 | 3 | 2 | 1.318 | 0.3182 | ns |
| 1D, SF1D | *MR1* mRNA | B2B | 6-FP | NT | B2B | 6-FP | MAIT | 3 | 3 | 2 | 1.466 | 0.2802 | ns |
| 1D, SF1D | *MR1* mRNA | B2B | UT | MAIT | B2B | 6-FP | MAIT | 3 | 3 | 2 | 0.8054 | 0.5051 | ns |
| 1D, SF1D | *MR1* mRNA | B2B | 5-OP | NT | B2B | 6-FP | NT | 3 | 3 | 2 | 0.5185 | 0.6557 | ns |
| 1D, SF1D | *MR1* mRNA | B2B | 5-OP | MAIT | B2B | 6-FP | MAIT | 3 | 3 | 2 | 5.239 | 0.0346 | * |
| 1E, SF1E | αMR1 gMFI | B2B | UT | NT | B2B | 5-OP | NT | 4 | 4 | 3 | 1.312 | 0.2808 | ns |
| 1E, SF1E | αMR1 gMFI | B2B | UT | NT | B2B | UT | MAIT | 4 | 4 | 3 | 0.9286 | 0.4216 | ns |
| 1E, SF1E | αMR1 gMFI | B2B | UT | NT | B2B | 5-OP | MAIT | 4 | 4 | 3 | 2.826 | 0.0664 | ns |
| 1E, SF1E | αMR1 gMFI | B2B | 5-OP | NT | B2B | UT | MAIT | 4 | 4 | 3 | 2.042 | 0.1338 | ns |
| 1E, SF1E | αMR1 gMFI | B2B | 5-OP | NT | B2B | 5-OP | MAIT | 4 | 4 | 3 | 3.166 | 0.0506 | ns |
| 1E, SF1E | αMR1 gMFI | B2B | UT | MAIT | B2B | 5-OP | MAIT | 4 | 4 | 3 | 3.218 | 0.0487 | * |
| 1E, SF1E | αMR1 gMFI | B2B | UT | NT | B2B | 6-FP | NT | 4 | 4 | 3 | 0.2705 | 0.8043 | ns |
| 1E, SF1E | αMR1 gMFI | B2B | UT | NT | B2B | 6-FP | MAIT | 4 | 4 | 3 | 0.5864 | 0.5988 | ns |
|  |  | Cell | Ag | T cell | Cell | Ag | T cell |  |  |  |  |  |  |
| 1E, SF1E | αMR1 gMFI | B2B | 6-FP | NT | B2B | UT | MAIT | 4 | 4 | 3 | 1.558 | 0.2171 | ns |
| 1E, SF1E | αMR1 gMFI | B2B | 6-FP | NT | B2B | 6-FP | MAIT | 4 | 4 | 3 | 1.349 | 0.2703 | ns |
| 1E, SF1E | αMR1 gMFI | B2B | UT | MAIT | B2B | 6-FP | MAIT | 4 | 4 | 3 | 0.03772 | 0.9723 | ns |
| 1E, SF1E | αMR1 gMFI | B2B | 5-OP | NT | B2B | 6-FP | NT | 4 | 4 | 3 | 2.235 | 0.1114 | ns |
| 1E, SF1E | αMR1 gMFI | B2B | 5-OP | MAIT | B2B | 6-FP | MAIT | 4 | 4 | 3 | 2.559 | 0.0833 | ns |

*Definition of abbreviations:*
Ag = antigen; df = degrees of freedom; statistic = absolute value of T statistic; AEC = airway epithelial cells; B2B = BEAS-2B cells; UI = uninfected control; *Sp* = *Streptococcus pneumoniae*; *Ms* = *Mycobacterium smegmatis*; UT = media treated control; 5-OP = 5-OP-RU; 6-FP = 6-formylpterin; NT = no T cell control; gMFI = geometric mean fluorescence intensity

We next asked whether microbial infection is required for this transcriptional increase or if the presence of MR1 ligand alone is sufficient. We treated BEAS-2B cells with either the stimulatory antigen 5-(2-oxopropylideneamino)-6-d-ribitylaminouracil (5-OP-RU) or the non-stimulatory ligand 6-formylpterin (6-FP) and measured expression of MR1. Neither 5-OP-RU nor 6-FP increased *MR1* expression alone (Figure 1D-E). In co-culture with MAIT cells, however, *MR1* mRNA expression increased when BEAS-2B cells were treated with 5-OP-RU, while treatment with 6-FP had no impact on *MR1* expression (Figure 1D-E, Supplemental Figure 1D-E). These data demonstrate that the upregulation of *MR1* mRNA expression requires activation of MAIT cells, and this may be stimulated by antigen presentation alone or bacterial infection.

### IFNγ stimulates MR1 expression and antigen presentation

Among the effector molecules produced by activated MAIT cells, IFNγ is well known to stimulate transcription of MHC Class I molecules^15–17^. We hypothesized *MR1* expression could be regulated through similar mechanisms, despite the differences in chromosomal location and gene arrangement from classical *HLA* genes. To test if IFNγ alone is sufficient to stimulate *MR1* expression, we treated primary AEC with recombinant human IFNγ. IFNγ treatment significantly increased *MR1* transcription (Figure 2A, left). As expected, IFNγ also increased expression of positive control *HLA-A* (Figure 2A, right). In BEAS-2B cells treated with IFNγ, we also observed significant increases in both *MR1* and *HLA-A* mRNA expression (Figure 2B).

**Figure 2.**
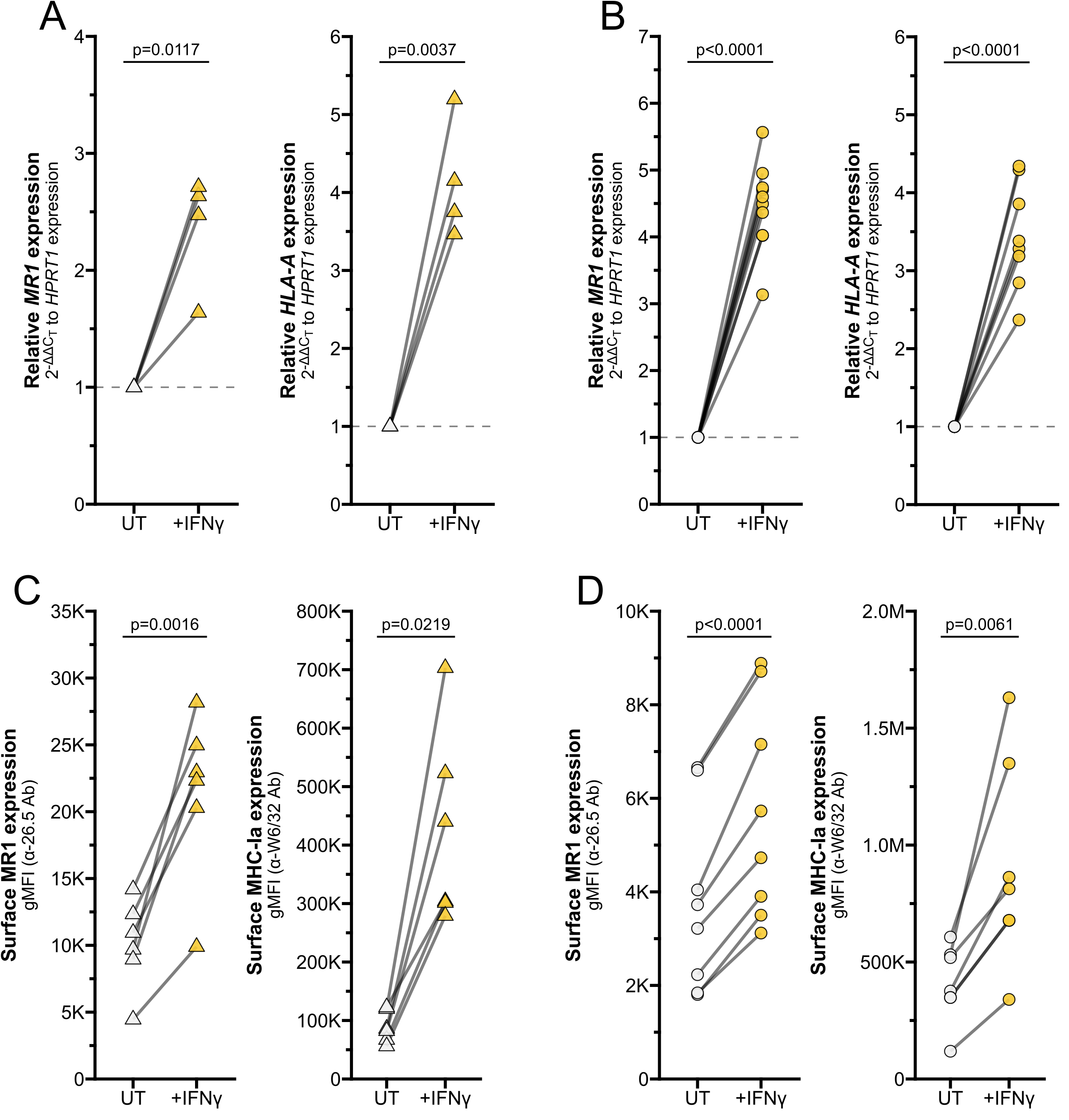
IFNγ induces MR1 expression and function. RT-qPCR of **(A)** primary human AECs or **(B)** BEAS-2B cells treated with media control (UT) or recombinant IFNγ for 12 hours. *MR1* (left) and *HLA-A* (right) expression were calculated relative to *HPRT1* expression and UT control, paired by individual donor or experimental replicate. Flow cytometry of **(C)** primary AECs or **(D)** BEAS-2B cells treated with recombinant IFNγ for 12 hours. gMFI of surface MR1 (left, α-26.5 Ab) and MHC-Ia (right, α-W6/32 Ab) are paired by individual donor or experimental replicate. Pairwise T tests were performed by donor **(A** and **C)** or experiment **(B** and **D)**.

To quantify MR1 protein expression, we measured surface MR1 expression by flow cytometry. We found that IFNγ treatment also significantly increased surface MR1 expression and control MHC-I expression in primary AEC and BEAS-2B cells (Figure 2C-D. This approach does not distinguish between 1) increased surface expression of MR1 proteins due to increased *MR1* transcription and translation or 2) increased translocation of existing MR1 molecules and stabilization on the cell surface. To determine if IFNγ signaling impacts post-transcriptional protein stability of MR1, we utilized BEAS-2B cells expressing MR1-GFP under a doxycycline-inducible promoter^11^. IFNγ treatment did not increase expression of *MR1* mRNA, total MR1-GFP gMFI, or surface MR1 in these doxycycline-treated cells (Figure 3A-C). As expected, IFNγ treatment increased MHC-Ia surface expression and 6-FP treatment induced significant stabilization of total MR1-GFP protein expression and surface translocation (Figure 3C-D). Together, these data indicate that increased surface MR1 in IFNγ-treated wildtype cells resulted from stimulation of *MR1* transcription rather than protein-level impacts.

**Figure 3.**
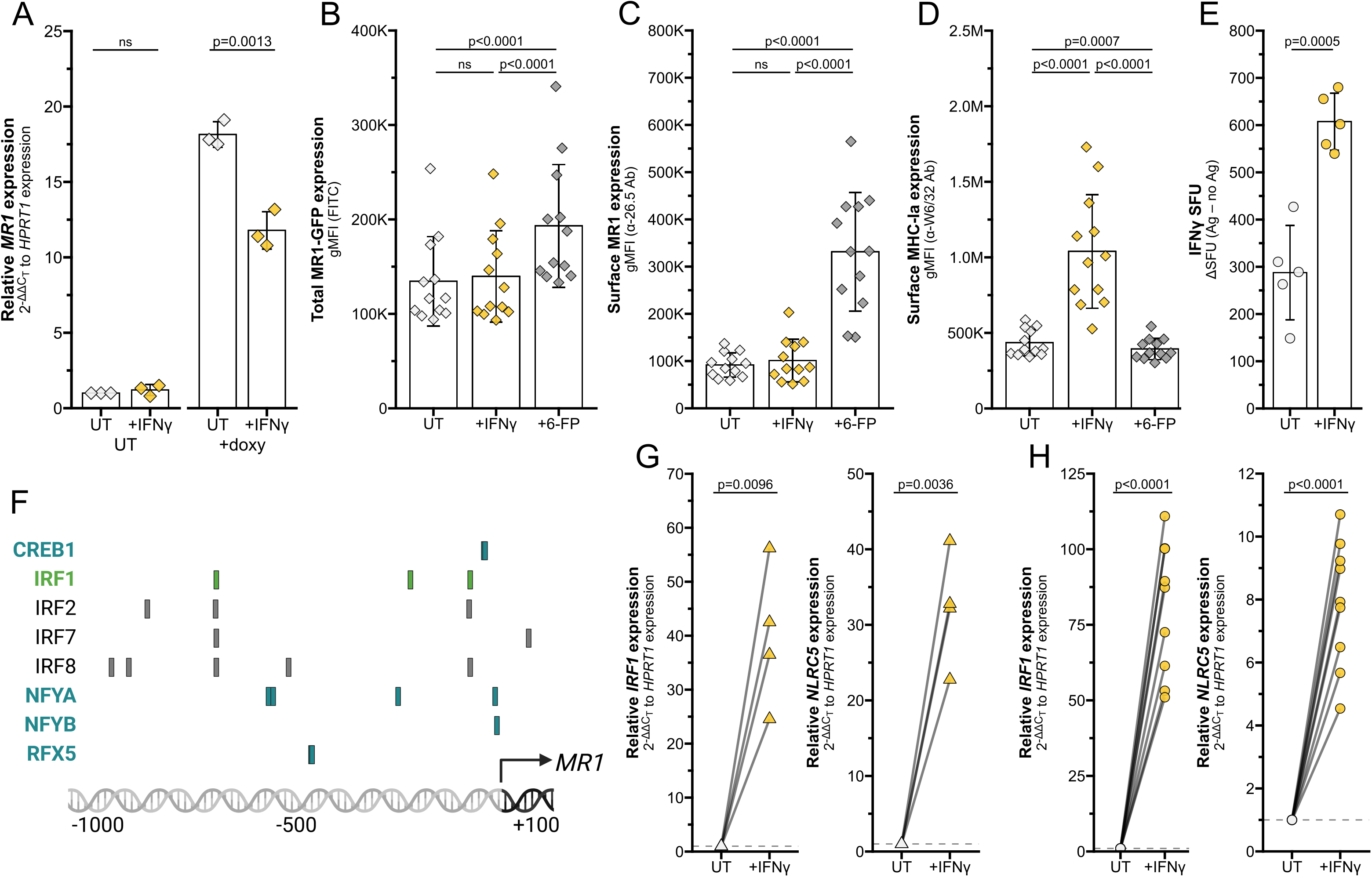
Transcriptional stimulation of MR1 by IFNγ. **(A-D)** BEAS-2B:doxMR1-GFP cells were treated with doxycycline, IFNγ, and/or 6-FP overnight. **(A)** *MR1* expression was calculated relative to *HPRT1* expression and UT control, paired by experimental replicate. gMFI of **(B)** MR1-GFP, **(C)** surface MR1 α-26.5 stain, and **(D)** surface MHC-Ia α-W6/32 stain. Data are experimental replicates. **(E)** ELISPOT of BEAS-2B cells treated with filtered *M. smegmatis* supernatant and MAIT cells. Data points are experimental replicates of no-antigen background-subtracted IFNγ spot-forming units (SFU). **(F)** Putative transcription factor binding sites were acquired through the Eukaryotic Promoter Database browser using the Search Motif Tool to perform on-the-fly scanning for transcription factor motifs using the FindM tool from the Signal Search Analysis (SSA) Server toolkit^28,97–99^. Highlighted proteins are involved in IRF1-(green) or NLRC5 enhanceosome-(blue) mediated *HLA* transcription. RT-qPCR of **(G)** primary human AECs or **(H)** BEAS-2B cells treated with recombinant IFNγ for 12 hours. *IRF1* (left) and *NLRC5* (right) expression were calculated relative to *HPRT1* expression and UT control, paired by individual donor or experimental replicate. Pairwise T tests were performed by experiment (A-E, H) or donor (G).

MR1 antigen presentation and MAIT cell responses are increased in MR1 over-expression systems^10,11,25^. We next investigated if the IFNγ-dependent increase in MR1 expression similarly enhanced MAIT cell responses to wildtype cells. BEAS-2B cells were treated with IFNγ for 12 hours, thoroughly washed to remove excess soluble IFNγ, then used as antigen-presenting cells in an IFNγ ELISPOT assay to quantify MAIT cell activation. Filtered *M. smegmatis* supernatant was used as the antigen source to avoid potential confounding impacts of IFNγ treatment on bacterial infection. MAIT cell responses to IFNγ pre-treated BEAS-2B cells were significantly increased compared to UT controls (Figure 3E). Therefore, IFNγ treatment is sufficient to stimulate *MR1* transcription, leading to increased protein expression and antigen presentation to MAIT cells.

### IFNγ stimulates MR1 transcription via transcription factor IRF1, not NLRC5

Using MHC-Ia transcription pathways as a starting point, we queried the JASPAR CORE 2018 Vertebrates database to determine if the *MR1* promoter contained binding motifs for known IFNγ-induced transcription factors^26–28^. We highlighted notable predicted elements on this region, including those common with IFNγ-mediated *HLA* transcription factor sites (Figure 3F). For example, we found putative binding motifs for IRF1 and members of the NLRC5 enhanceosome complex^18,19,29–31^. This targeted search suggested that IRF1 and NLRC5 could be of interest in IFNγ-mediated *MR1* transcription.

We first validated that IFNγ signaling induces *IRF1* and *NLRC5* mRNA expression in our cells. Transcripts of both these genes were significantly increased by IFNγ treatment of both primary human AEC and BEAS-2B cells (Figure 3G-H).

To determine if IRF1 or NLRC5 are required for the IFNγ-mediated increase in *MR1* transcription, we first used siRNA to knock down the genes separately or together in BEAS-2B cells. IRF1 KD alone significantly decreased IFNγ-stimulated *MR1* mRNA expression compared to missense controls (Figure 4A, Table 2). Although IRF1 siRNA significantly reduced *IRF1* expression, we were unable to sufficiently knock down *NLRC5* expression by siRNA (Supplemental Figure 2A-B). Therefore, we generated monoclonal NLRC5^−/−^ BEAS-2B cell lines by CRISPR/Cas9. Loss of NLRC5 did not lead to any significant impact to IFNγ-induced *MR1* mRNA expression (Figure 4B). We did observe a decrease in *HLA-A* expression with NLRC5 knockout, although not significant (Figure 4C).

**Figure 4.**
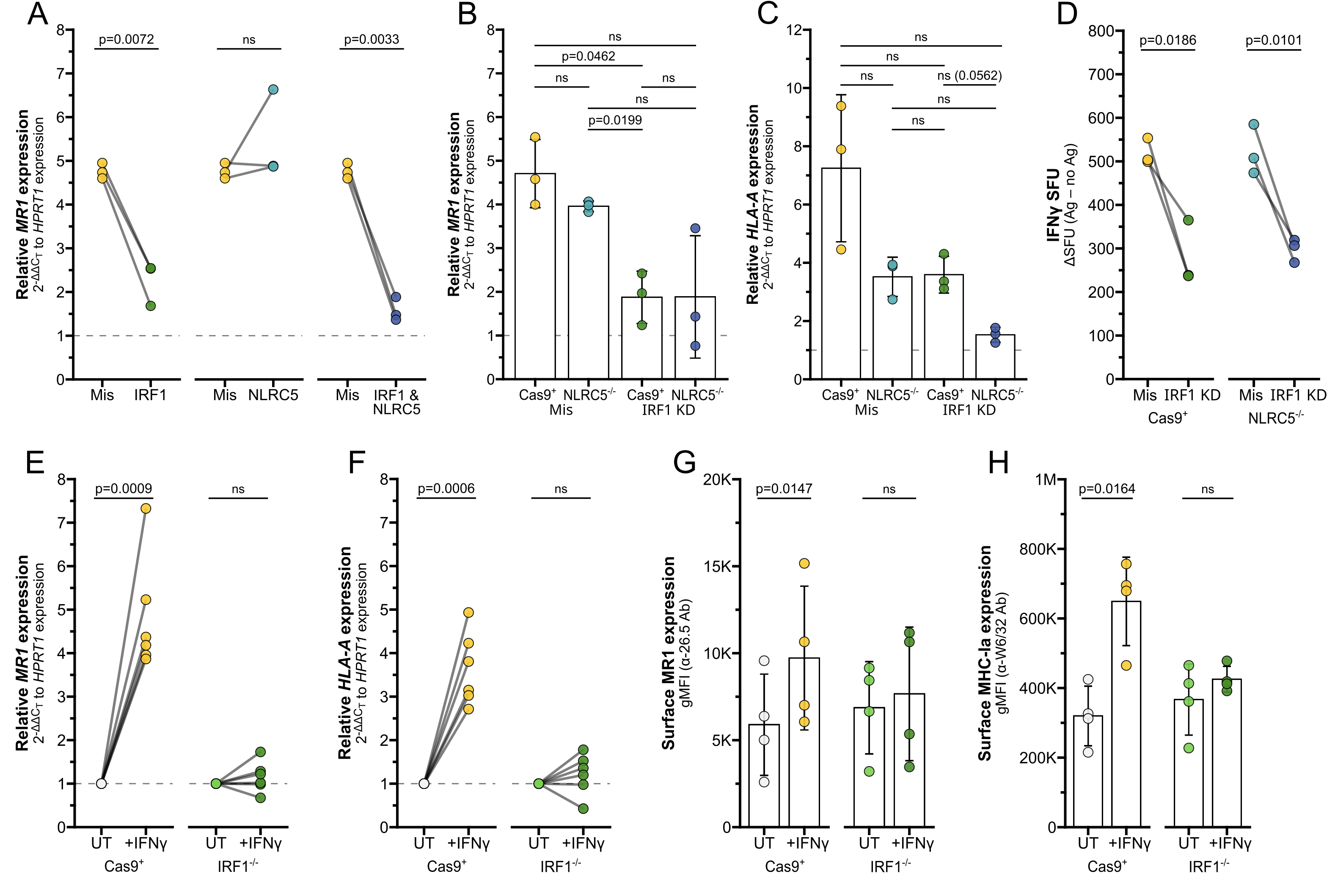
IRF1 mediates IFNγ-induced MR1 transcription. **(A)** RT-qPCR of BEAS-2B cells treated with IRF1, NLRC5, and/or missense siRNA as labeled for 36 hours, then incubated with IFNγ for 12 hours. *MR1* expression was calculated relative to *HPRT1* expression and missense UT control, paired by experimental replicate. **(B-C)** RT-qPCR of Cas9^+^ or NLRC5^−/−^ clone #1 BEAS-2B cells treated with IRF1 or missense siRNA for 36 hours, then incubated with IFNγ for 12 hours. **(B)** *MR1* and **(C)** *HLA-A* expression were calculated relative to *HPRT1* expression and Cas9^+^ or NLRC5^−/−^ clone #1 missense UT controls, paired by experimental replicate. **(D)** Cells in (B-C) were used as antigen-presenting cells in ELISPOT assay, with filtered *M. smegmatis* supernatant as the antigen source. Data points are experimental replicates of Cas9^+^ or NLRC5^−/−^ clone #1 missense control no-antigen background-subtracted IFNγ SFU. **(E-F)** RT-qPCR of Cas9^+^ or IRF1^−/−^ clone #2 BEAS-2B cells treated with IFNγ for 12 hours. **(E)** *MR1* and **(F)** *HLA-A* expression were calculated relative to *HPRT1* expression and Cas9^+^ or IRF1^−/−^ clone #2 UT controls, paired by experimental replicate. **(G-H)** Flow cytometry of Cas9^+^ or IRF1^−/−^ clone #1 BEAS-2B cells treated with IFNγ for 12 hours. gMFI of **(G)** surface MR1 (α-26.5 Ab) and **(H)** MHC-Ia (α-W6/32 Ab) are paired by experimental replicate. Statistical analyses are in Table 2.

**Table 2.**
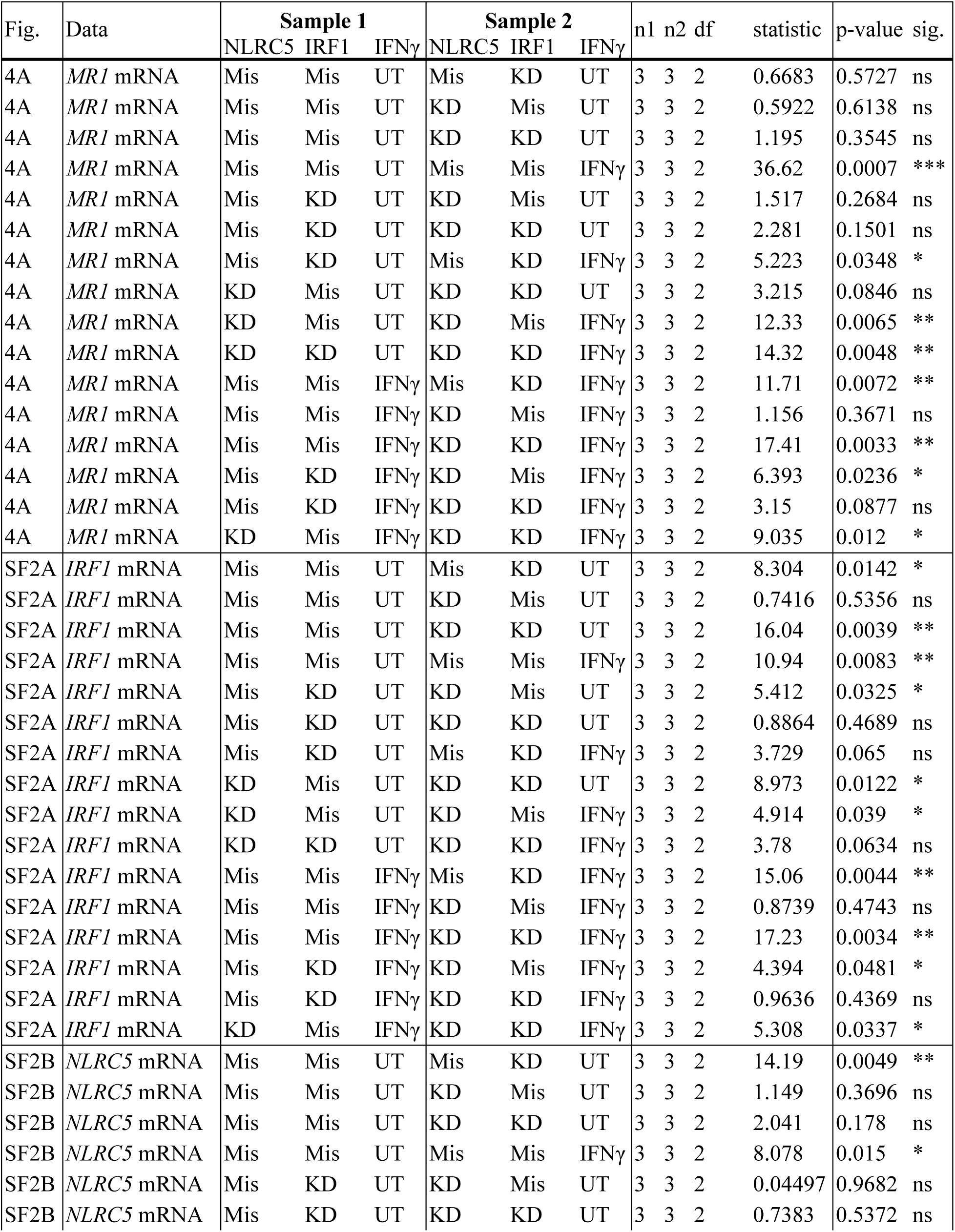
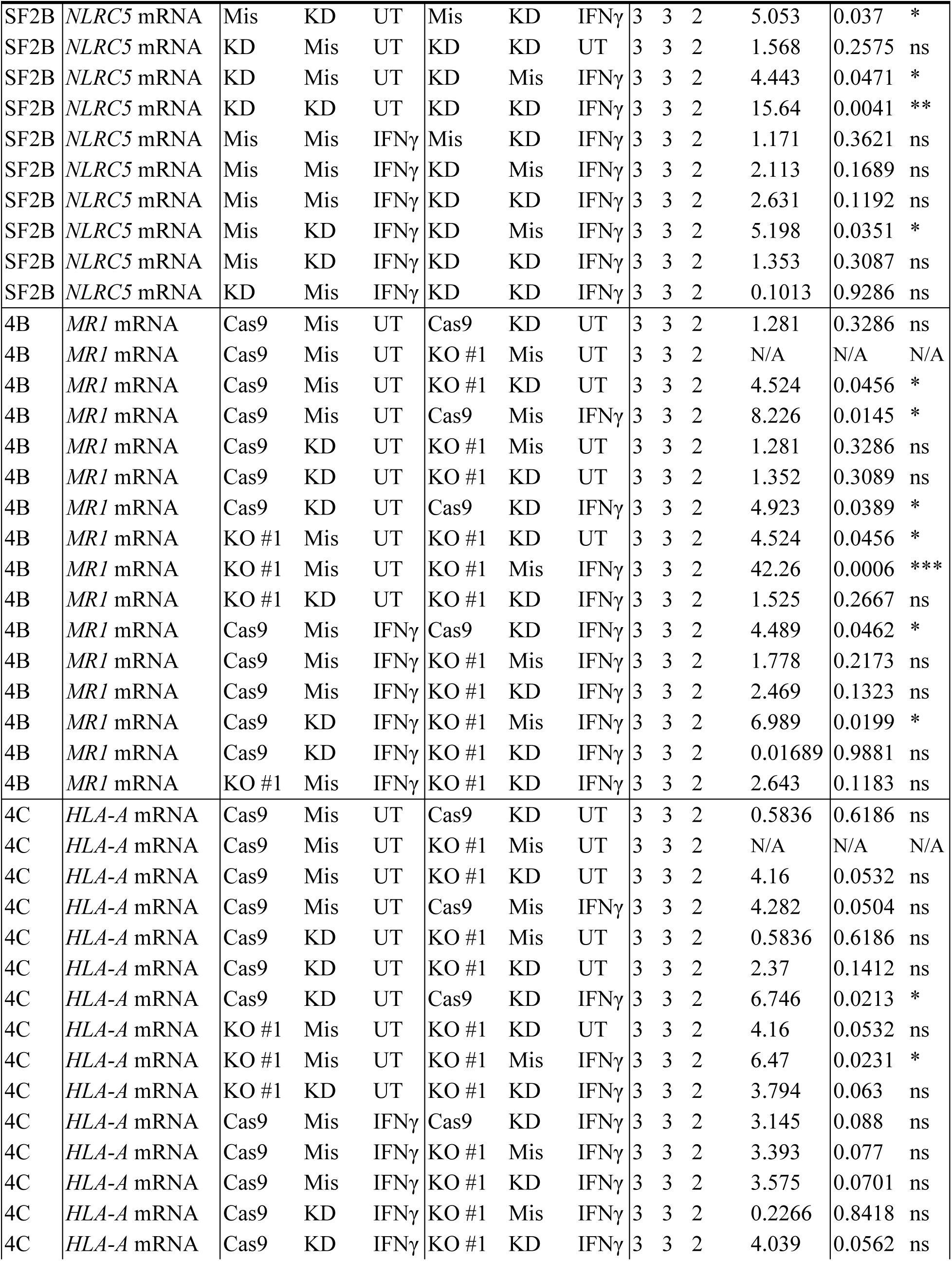
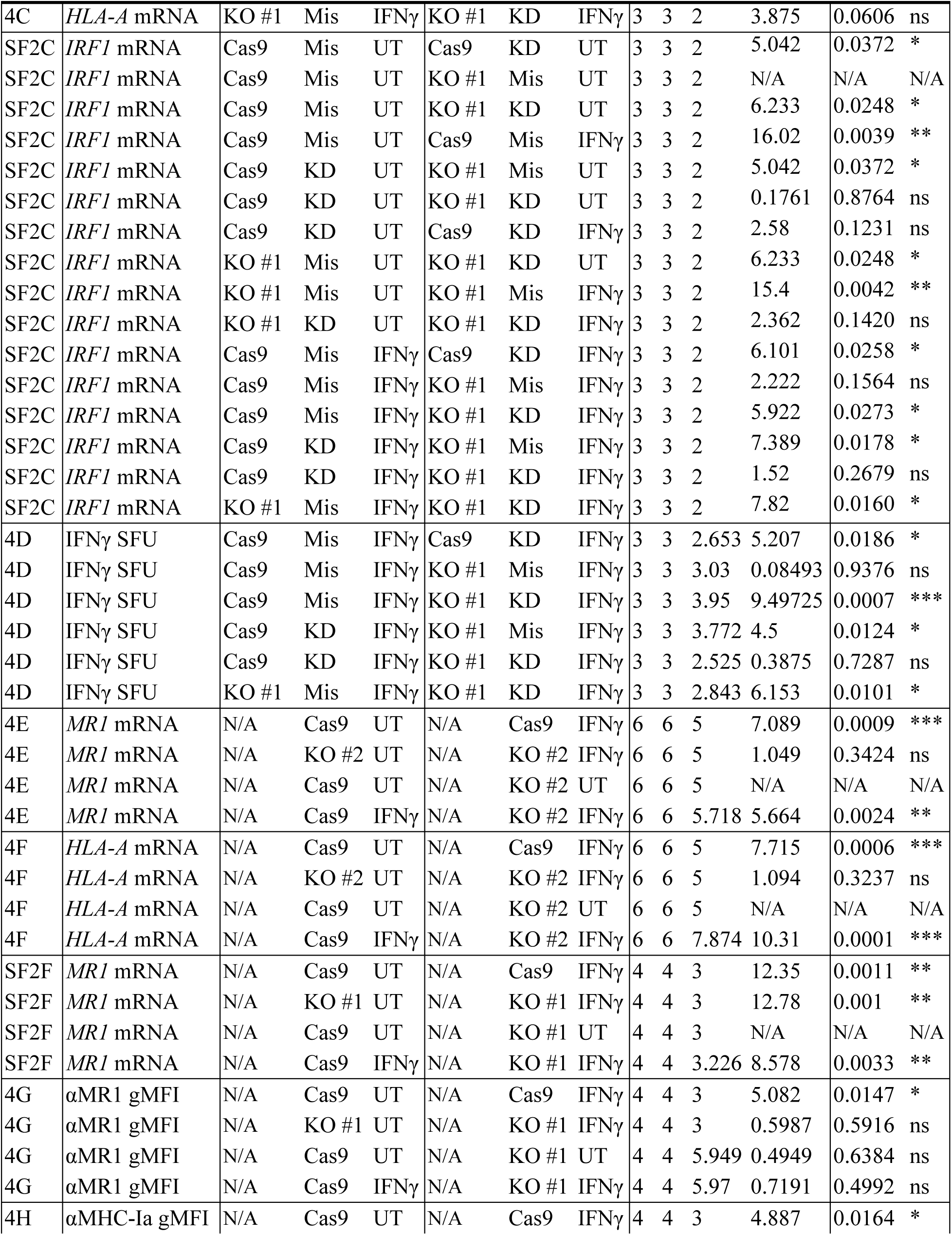
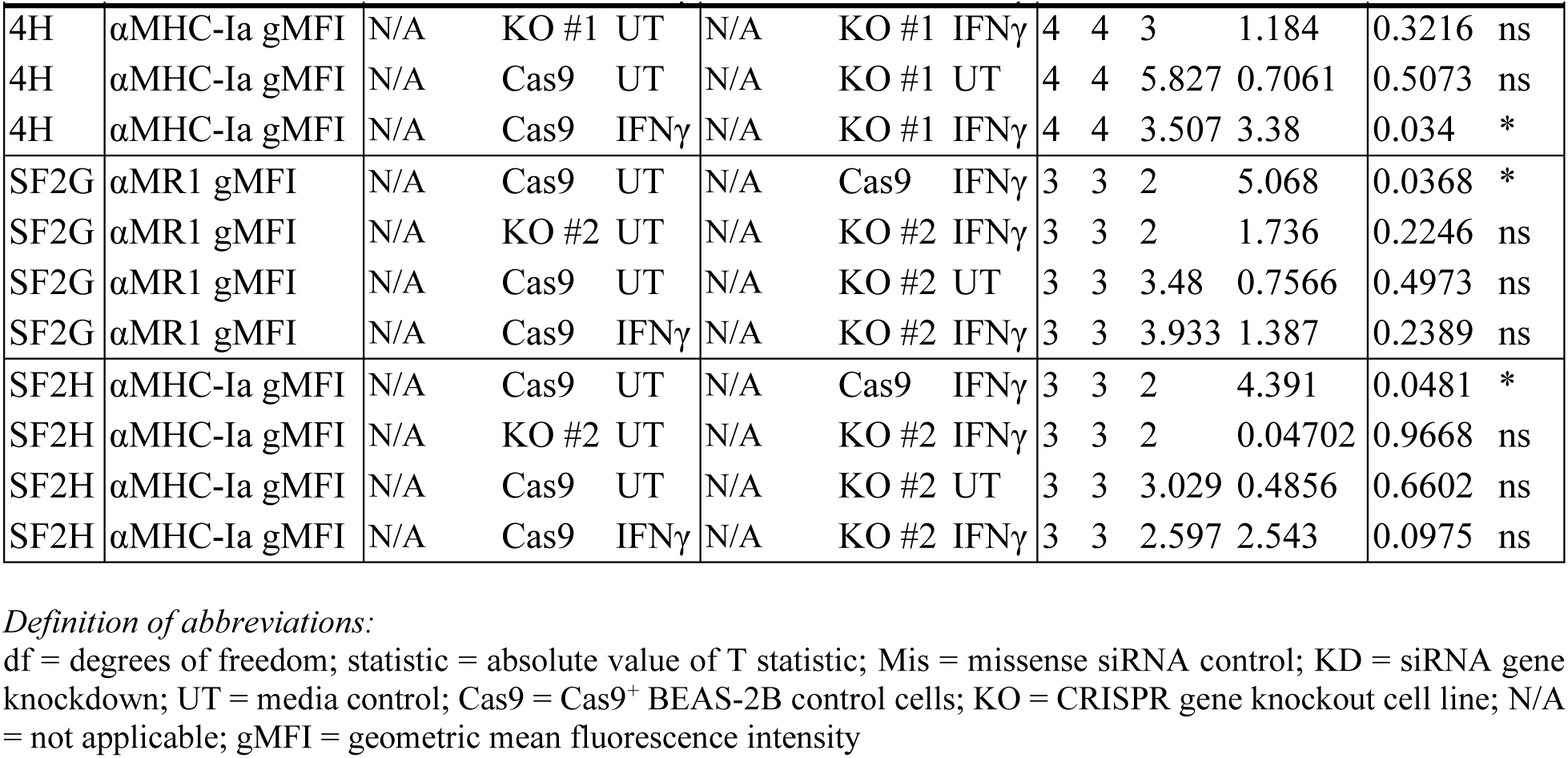
Statistics associated with Figure 4 and Supplemental Figure 2.

We used siRNA to silence IRF1 expression in the Cas9^+^ and NLRC5^−/−^ cells (Supplemental Figure 2C). *MR1* expression was significantly impacted by IRF1 knockdown in Cas9^+^ control cells (Figure 4B), agreeing with the previous siRNA results in wildtype BEAS-2b cells. A trend in reduced *MR1* expression was also observed in two clones of NLRC5^−/−^ cells treated with IRF1 siRNA (Figure 4B, Supplemental Figure 2D). Expression of surface MR1 proteins in IFNγ-treated cells was similarly decreased with IRF1 silencing and unaffected by NLRC5 knockout (Supplemental Figure 2E). Finally, we used ELISPOT assays to quantify if loss of NLRC5 and/or IRF1 would impact the IFNγ-stimulated boost in MR1 antigen presentation to MAIT cells. IRF1 siRNA knockdown significantly reduced MAIT cell responses to both Cas9^+^ cells and NLRC5^−/−^ cells (Figure 4D). The missense-treated NLRC5^−/−^ cells stimulated similar MAIT cell activity as the Cas9^+^ control cells in response to IFNγ (Figure 4D, Table 2). Together, these results indicate that IRF1 expression is required for IFNγ stimulation of MR1 expression and antigen presentation function, while NLRC5 does not appear to impact this pathway.

We next used CRISPR/Cas9 knockout to generate monoclonal IRF1^−/−^ BEAS-2B cell lines to confirm this finding. IFNγ-mediated stimulation of *MR1* mRNA and MR1 surface expression was impaired in IRF1^−/−^ cells compared to Cas9^+^ cells (Figure 4E,G, Supplemental Figure 2F-G). Expression of *HLA-A* mRNA or MHC-Ia surface proteins were likewise impaired in the IRF1^−/−^ cells following IFNγ treatment (Figure 4F,H; Supplemental Figure 2G, right). Together the siRNA knockdown and CRISPR knockout results indicate that IRF1 mediates the IFNγ signaling pathway leading to MR1 transcription, surface expression, and antigen presentation, likely through mechanisms independent of the NLRC5 enhanceosome.

### MAIT cells produce IFNγ sufficient to induce MR1 transcription pathways

To validate this mechanism in a physiologically relevant system, we returned to our co-culture experiments. We first demonstrated that co-culture of infected AEC with MAIT cells led to upregulation of IFNγ-stimulated pathways by staining phosphorylated STAT1 (pSTAT1). Ligation of the IFNGR activates Janus kinase (JAK) dimer 1/2, which in turn phosphorylates STAT1^17,32^. As expected, pSTAT1 staining was significantly increased in AEC infected with *S. pneumoniae* and co-cultured with MAIT cells, along with AEC treated with recombinant IFNγ (Figure 5A-B, Supplemental Figure 3A, Table 3).

**Figure 5.**
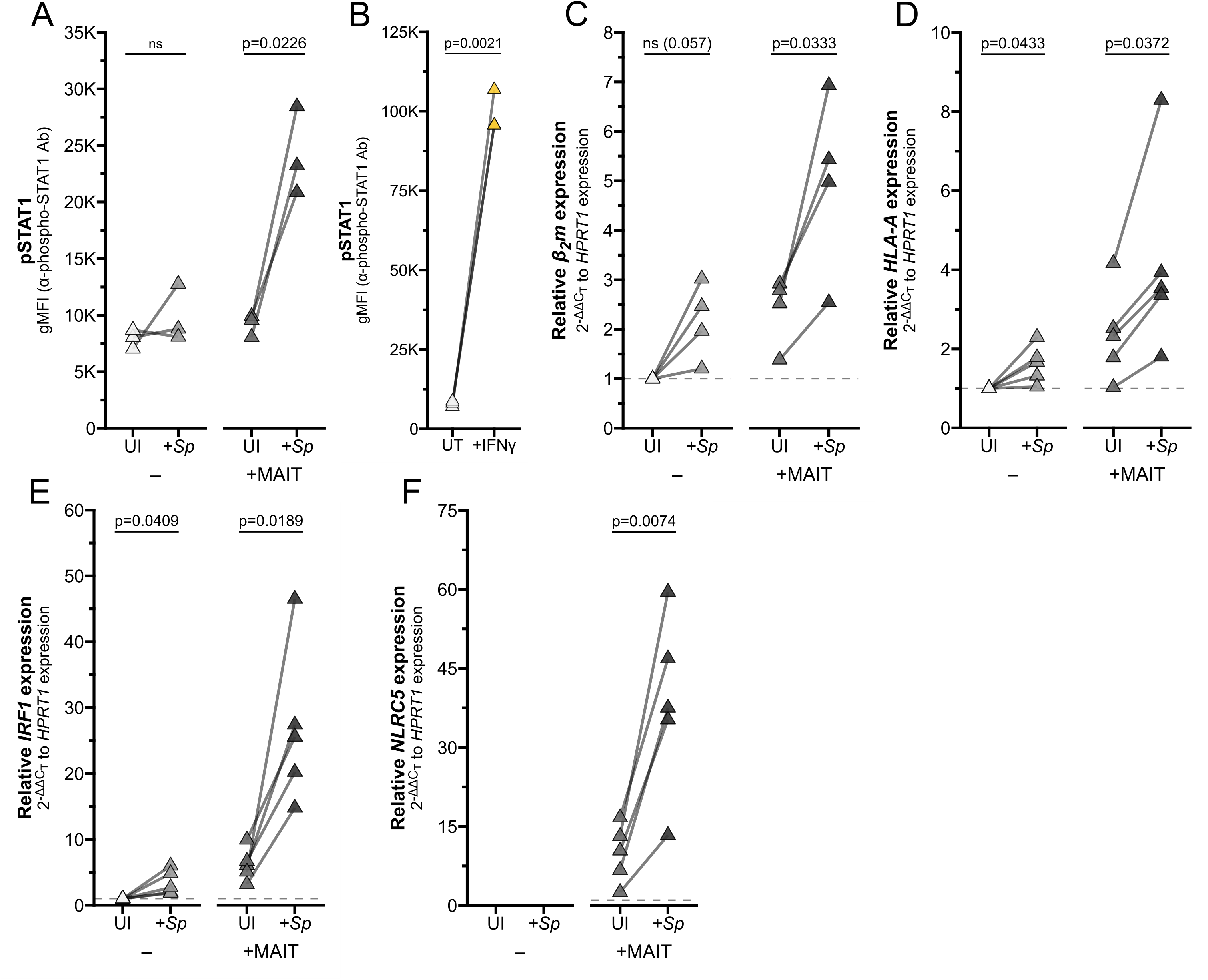
Reciprocal IFNγ signaling in *Sp*-infected AEC co-cultured with MAIT cells. **(A-B)** Flow cytometry of **(A)** primary human AECs infected with *S. pneumoniae* (*Sp*) for one hour and incubated overnight with MAIT cell clone (n=3), or **(B)** AECs treated with IFNγ for 12 hours (n=3). gMFI of stained pSTAT1 expression is paired by individual donor. **(C-F)** RT-qPCR of primary human AECs infected with *S. pneumoniae* (*Sp*) for one hour and incubated overnight with MAIT cell clone. Expression of **(C)** *β2m,* **(D)** *HLA-A* **(E)** *IRF1*, and **(F)** *NLRC5* were calculated relative to *HPRT1* expression and UI-control, paired by individual donor (n=4 (C) or n=5 (D-F) donors). Pairwise statistical analyses are in Table 3.

**Table 3.** Statistics associated with Figure 5, Figure 6, and Supplemental Figure 3.

| Figure | Data | Sample 1 |  |  | Sample 2 |  |  | n1 | n2 | df | statistic | p-value | sig. |
| --- | --- | --- | --- | --- | --- | --- | --- | --- | --- | --- | --- | --- | --- |
|  |  | Cell | Ag | T cell | Cell | Ag | T cell |  |  |  |  |  |  |
| 5A, SF3A | pSTAT1 gMFI | AEC | UI | NT | AEC | *Sp* | NT | 3 | 3 | 2 | 1.018 | 0.4159 | ns |
| 5A, SF3A | pSTAT1 gMFI | AEC | UI | NT | AEC | UI | MAIT | 3 | 3 | 2 | 1.472 | 0.2788 | ns |
| 5A, SF3A | pSTAT1 gMFI | AEC | UI | NT | AEC | *Sp* | MAIT | 3 | 3 | 2 | 9.043 | 0.012 | * |
| 5A, SF3A | pSTAT1 gMFI | AEC | *Sp* | NT | AEC | UI | MAIT | 3 | 3 | 2 | 0.5673 | 0.6277 | ns |
| 5A, SF3A | pSTAT1 gMFI | AEC | *Sp* | NT | AEC | *Sp* | MAIT | 3 | 3 | 2 | 4.031 | 0.0564 | ns |
| 5A, SF3A | pSTAT1 gMFI | AEC | UI | MAIT | AEC | *Sp* | MAIT | 3 | 3 | 2 | 6.541 | 0.0226 | * |
| 5C, SF3B | *B2m* mRNA | AEC | UI | NT | AEC | *Sp* | NT | 4 | 4 | 3 | 3.014 | 0.057 | ns |
| 5C, SF3B | *B2m* mRNA | AEC | UI | NT | AEC | UI | MAIT | 4 | 4 | 3 | 4.013 | 0.0278 | * |
| 5C, SF3B | *B2m* mRNA | AEC | UI | NT | AEC | *Sp* | MAIT | 4 | 4 | 3 | 4.361 | 0.0223 | * |
| 5C, SF3B | *B2m* mRNA | AEC | *Sp* | NT | AEC | UI | MAIT | 4 | 4 | 3 | 0.9405 | 0.4164 | ns |
| 5C, SF3B | *B2m* mRNA | AEC | *Sp* | NT | AEC | *Sp* | MAIT | 4 | 4 | 3 | 4.302 | 0.0231 | * |
| 5C, SF3B | *B2m* mRNA | AEC | UI | MAIT | AEC | *Sp* | MAIT | 4 | 4 | 3 | 3.741 | 0.0333 | * |
| 5D, SF3D | *HLA-A* mRNA | AEC | UI | NT | AEC | *Sp* | NT | 5 | 5 | 4 | 2.918 | 0.0433 | * |
| 5D, SF3D | *HLA-A* mRNA | AEC | UI | NT | AEC | UI | MAIT | 5 | 5 | 4 | 2.629 | 0.0582 | ns |
| 5D, SF3D | *HLA-A* mRNA | AEC | UI | NT | AEC | *Sp* | MAIT | 5 | 5 | 4 | 2.925 | 0.043 | * |
| 5D, SF3D | *HLA-A* mRNA | AEC | *Sp* | NT | AEC | UI | MAIT | 5 | 5 | 4 | 1.346 | 0.2496 | ns |
| 5D, SF3D | *HLA-A* mRNA | AEC | *Sp* | NT | AEC | *Sp* | MAIT | 5 | 5 | 4 | 2.268 | 0.0859 | ns |
| 5D, SF3D | *HLA-A* mRNA | AEC | UI | MAIT | AEC | *Sp* | MAIT | 5 | 5 | 4 | 3.072 | 0.0372 | * |
| 5E, SF3G | *IRF1* mRNA | AEC | UI | NT | AEC | *Sp* | NT | 5 | 5 | 4 | 2.977 | 0.0409 | * |
| 5E, SF3G | *IRF1* mRNA | AEC | UI | NT | AEC | UI | MAIT | 5 | 5 | 4 | 4.702 | 0.0093 | ** |
| 5E, SF3G | *IRF1* mRNA | AEC | UI | NT | AEC | *Sp* | MAIT | 5 | 5 | 4 | 4.822 | 0.0085 | ** |
| 5E, SF3G | *IRF1* mRNA | AEC | *Sp* | NT | AEC | UI | MAIT | 5 | 5 | 4 | 3.392 | 0.0275 | * |
| 5E, SF3G | *IRF1* mRNA | AEC | *Sp* | NT | AEC | *Sp* | MAIT | 5 | 5 | 4 | 4.703 | 0.0093 | ** |
| 5E, SF3G | *IRF1* mRNA | AEC | UI | MAIT | AEC | *Sp* | MAIT | 5 | 5 | 4 | 3.813 | 0.0189 | * |
| 5F, SF3C | *NLRC5* mRNA | AEC | UI | NT | AEC | *Sp* | NT | 5 | 5 | 4 | 2.084 | 0.1055 | ns |
| 5F, SF3C | *NLRC5* mRNA | AEC | UI | NT | AEC | UI | MAIT | 5 | 5 | 4 | 3.605 | 0.0227 | * |
| 5F, SF3C | *NLRC5* mRNA | AEC | UI | NT | AEC | *Sp* | MAIT | 5 | 5 | 4 | 4.933 | 0.0079 | ** |
| 5F, SF3C | *NLRC5* mRNA | AEC | *Sp* | NT | AEC | UI | MAIT | 5 | 5 | 4 | 0.6506 | 0.5508 | ns |
| 5F, SF3C | *NLRC5* mRNA | AEC | *Sp* | NT | AEC | *Sp* | MAIT | 5 | 5 | 4 | 3.829 | 0.0186 | * |
| 5F, SF3C | *NLRC5* mRNA | AEC | UI | MAIT | AEC | *Sp* | MAIT | 5 | 5 | 4 | 5.008 | 0.0074 | ** |
| 6A, SF3E | *HLA-A* mRNA | B2B | UI | NT | B2B | *Ms* | NT | 3 | 3 | 2 | 0.09115 | 0.9357 | ns |
| 6A, SF3E | *HLA-A* mRNA | B2B | UI | NT | B2B | UI | MAIT | 3 | 3 | 2 | 2.963 | 0.0975 | ns |
| 6A, SF3E | *HLA-A* mRNA | B2B | UI | NT | B2B | *Ms* | MAIT | 3 | 3 | 2 | 5.839 | 0.0281 | * |
| 6A, SF3E | *HLA-A* mRNA | B2B | *Ms* | NT | B2B | UI | MAIT | 3 | 3 | 2 | 3.015 | 0.0946 | ns |
| 6A, SF3E | *HLA-A* mRNA | B2B | *Ms* | NT | B2B | *Ms* | MAIT | 3 | 3 | 2 | 5.658 | 0.0298 | * |
| 6A, SF3E | *HLA-A* mRNA | B2B | UI | MAIT | B2B | *Ms* | MAIT | 3 | 3 | 2 | 4.79 | 0.0409 | * |
| 6B, SF3F | *HLA-A* mRNA | B2B | UT | NT | B2B | 5-OP | NT | 3 | 3 | 2 | 0.3951 | 0.7309 | ns |
|  |  | Cell | Ag | T cell | Cell | Ag | T cell |  |  |  |  |  |  |
| 6B, SF3F | *HLA-A* mRNA | B2B | UT | NT | B2B | UT | MAIT | 3 | 3 | 2 | 2.963 | 0.0975 | ns |
| 6B, SF3F | *HLA-A* mRNA | B2B | UT | NT | B2B | 5-OP | MAIT | 3 | 3 | 2 | 11.72 | 0.0072 | ** |
| 6B, SF3F | *HLA-A* mRNA | B2B | 5-OP | NT | B2B | UT | MAIT | 3 | 3 | 2 | 3.413 | 0.0762 | ns |
| 6B, SF3F | *HLA-A* mRNA | B2B | 5-OP | NT | B2B | 5-OP | MAIT | 3 | 3 | 2 | 10.2 | 0.0095 | ** |
| 6B, SF3F | *HLA-A* mRNA | B2B | UT | MAIT | B2B | 5-OP | MAIT | 3 | 3 | 2 | 10.94 | 0.0083 | ** |
| 6B, SF3F | *HLA-A* mRNA | B2B | UT | NT | B2B | 6-FP | NT | 3 | 3 | 2 | 0.1102 | 0.9223 | ns |
| 6B, SF3F | *HLA-A* mRNA | B2B | UT | NT | B2B | 6-FP | MAIT | 3 | 3 | 2 | 2.58 | 0.1231 | ns |
| 6B, SF3F | *HLA-A* mRNA | B2B | 6-FP | NT | B2B | UT | MAIT | 3 | 3 | 2 | 2.7 | 0.1142 | ns |
| 6B, SF3F | *HLA-A* mRNA | B2B | 6-FP | NT | B2B | 6-FP | MAIT | 3 | 3 | 2 | 2.416 | 0.1370 | ns |
| 6B, SF3F | *HLA-A* mRNA | B2B | UT | MAIT | B2B | 6-FP | MAIT | 3 | 3 | 2 | 0.3708 | 0.7464 | ns |
| 6B, SF3F | *HLA-A* mRNA | B2B | 5-OP | NT | B2B | 6-FP | NT | 3 | 3 | 2 | 0.4784 | 0.6796 | ns |
| 6C, SF3H | *IRF1* mRNA | B2B | UI | NT | B2B | *Ms* | NT | 3 | 3 | 2 | 0.5695 | 0.6264 | ns |
| 6C, SF3H | *IRF1* mRNA | B2B | UI | NT | B2B | UI | MAIT | 3 | 3 | 2 | 2.556 | 0.125 | ns |
| 6C, SF3H | *IRF1* mRNA | B2B | UI | NT | B2B | *Ms* | MAIT | 3 | 3 | 2 | 7.917 | 0.0156 | * |
| 6C, SF3H | *IRF1* mRNA | B2B | *Ms* | NT | B2B | UI | MAIT | 3 | 3 | 2 | 2.662 | 0.1169 | ns |
| 6C, SF3H | *IRF1* mRNA | B2B | *Ms* | NT | B2B | *Ms* | MAIT | 3 | 3 | 2 | 7.927 | 0.0155 | * |
| 6C, SF3H | *IRF1* mRNA | B2B | UI | MAIT | B2B | *Ms* | MAIT | 3 | 3 | 2 | 7.303 | 0.0182 | * |
| 6D, SF3I | *IRF1* mRNA | B2B | UT | NT | B2B | 5-OP | NT | 3 | 3 | 2 | 1.442 | 0.2860 | ns |
| 6D, SF3I | *IRF1* mRNA | B2B | UT | NT | B2B | UT | MAIT | 3 | 3 | 2 | 2.556 | 0.1250 | ns |
| 6D, SF3I | *IRF1* mRNA | B2B | UT | NT | B2B | 5-OP | MAIT | 3 | 3 | 2 | 7.652 | 0.0167 | * |
| 6D, SF3I | *IRF1* mRNA | B2B | 5-OP | NT | B2B | UT | MAIT | 3 | 3 | 2 | 2.693 | 0.1147 | ns |
| 6D, SF3I | *IRF1* mRNA | B2B | 5-OP | NT | B2B | 5-OP | MAIT | 3 | 3 | 2 | 7.67 | 0.0166 | * |
| 6D, SF3I | *IRF1* mRNA | B2B | UT | MAIT | B2B | 5-OP | MAIT | 3 | 3 | 2 | 8.056 | 0.0151 | * |
| 6D, SF3I | *IRF1* mRNA | B2B | UT | NT | B2B | 6-FP | NT | 3 | 3 | 2 | 0.212 | 0.8518 | ns |
| 6D, SF3I | *IRF1* mRNA | B2B | UT | NT | B2B | 6-FP | MAIT | 3 | 3 | 2 | 2.399 | 0.1385 | ns |
| 6D, SF3I | *IRF1* mRNA | B2B | 6-FP | NT | B2B | UT | MAIT | 3 | 3 | 2 | 2.539 | 0.1263 | ns |
| 6D, SF3I | *IRF1* mRNA | B2B | 6-FP | NT | B2B | 6-FP | MAIT | 3 | 3 | 2 | 2.392 | 0.1392 | ns |
| 6D, SF3I | *IRF1* mRNA | B2B | UT | MAIT | B2B | 6-FP | MAIT | 3 | 3 | 2 | 3.075 | 0.0915 | ns |
| 6D, SF3I | *IRF1* mRNA | B2B | 5-OP | NT | B2B | 6-FP | NT | 3 | 3 | 2 | 1.361 | 0.3065 | ns |

*Definition of abbreviations:*
Ag = antigen; df = degrees of freedom; statistic = absolute value of T statistic; AEC = airway epithelial cells; B2B = BEAS-2B cells; UI = uninfected control; *Sp* = *Streptococcus pneumoniae*; *Ms* = *Mycobacterium smegmatis*; UT = media treated control; 5-OP = 5-OP-RU; 6-FP = 6-formylpterin; NT = no T cell control

To confirm that activation of MAIT cells in co-culture is sufficient to drive IFNγ signaling, we next assessed expression of IFNγ-stimulated genes. Co-culture of *Sp*-infected primary AEC with MAIT cells significantly induced expression of *HLA-A, B2m, IRF1,* and *NLRC5* (Figure 5C-F; Supplemental Figure 3B-D,G; Table 3). We observed increased expression of these genes in AEC incubated with either *Sp* or MAIT cells alone, which may indicate the contribution of other inflammatory signaling pathways in AEC. However, the combined co-culture induced significantly greater expression for almost all genes, pointing to the role of Ag-induced MAIT cell activation in driving inflammatory gene expression.

BEAS-2B infected with *Ms* significantly induced expression of *HLA-A* and *IRF1* only in combination with MAIT cell co-culture (Figure 6A,C; Supplemental Figure 3E,H; Table 3). We next used 5-OP-RU as the antigen source to confirm that MR1 antigen presentation alone is sufficient to stimulate MAIT cell IFNγ production and subsequent inflammatory gene expression, absent other microbial stimuli. As expected, *HLA-A* and *IRF1* expression were significantly increased in 5-OP-RU-treated BEAS-2B cells when MAIT cells were present, but not stimulated by treatment with 5-OP-RU alone or MAIT cell co-culture with untreated BEAS-2B cells (Figure 6B,D; Supplemental Figure 3F,I; Table 3). Non-stimulatory presentation of 6-FP ligands failed to significantly induce either gene alone or in combination with MAIT cells. Together, these results indicate that activated MAIT cells produce sufficient IFNγ to stimulate expression of downstream genes.

**Figure 6.**
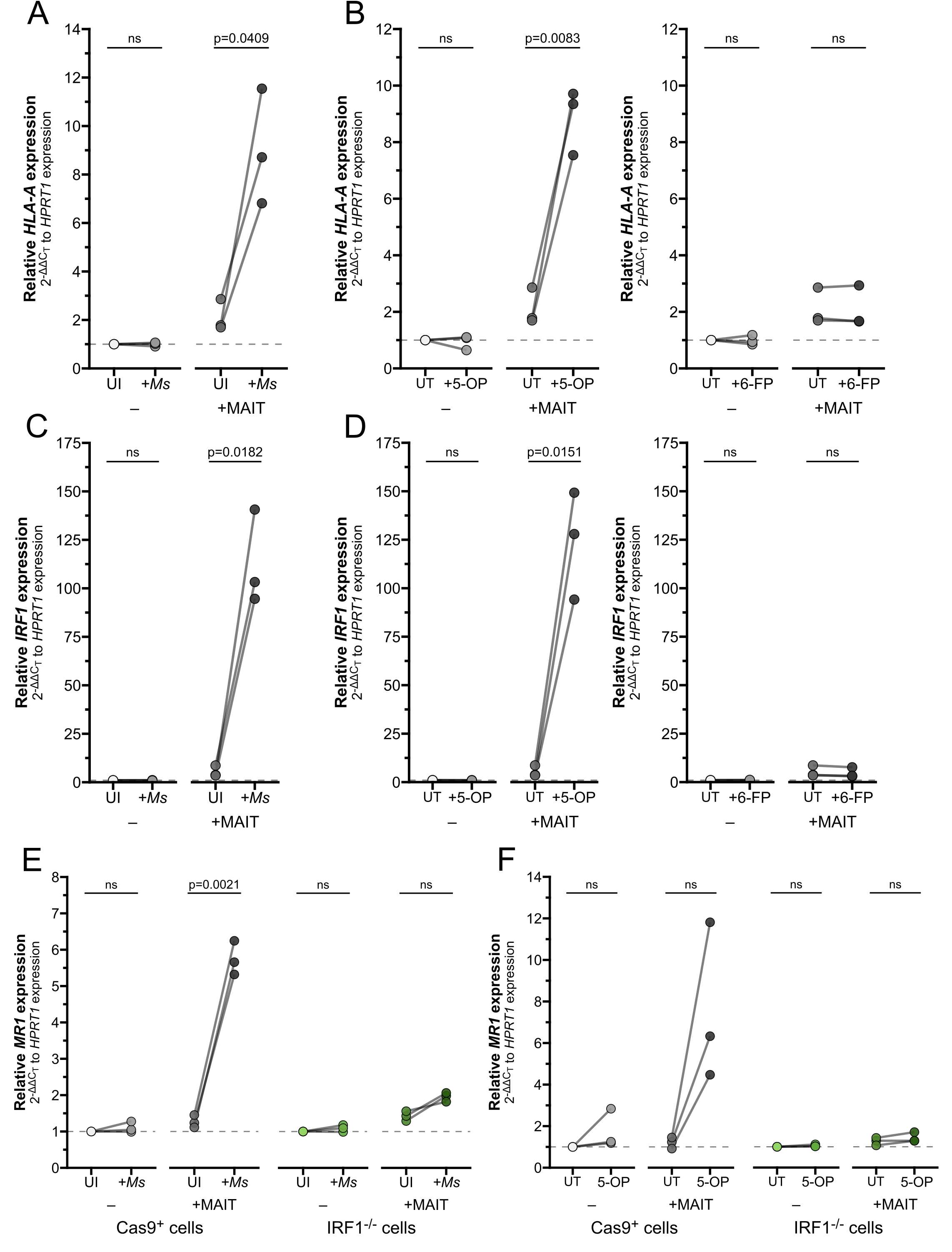
IFNγ produced by activated MAIT cells drives IRF1-dependent *MR1* transcription. **(A-D)** RT-qPCR of wildtype BEAS-2B cells treated as indicated below. Gene expression was calculated relative to *HPRT1* expression and UI- or UT-controls, paired by experiment. **(A)** *HLA-A* and **(C)** *IRF1* expression of BEAS-2B cells infected with *M. smegmatis* (*Ms*) for one hour and incubated overnight with MAIT cell clone. **(B)** *HLA-A* and **(D)** *IRF1* expression of BEAS-2B cells treated with 5-OP-RU (left, “5-OP”) or 6-FP (right) for one hour and incubated overnight with MAIT cell clone. **(E-F)** RT-qPCR of Cas9^+^ or IRF1^−/−^ clone #1 BEAS-2B cells **(E)** infected with *M. smegmatis* or **(F)** treated with 5-OP-RU for one hour, then incubated overnight with MAIT cell clone. *MR1* expression was calculated relative to *HPRT1* expression and Cas9^+^ or IRF1^−/−^ UT-controls, paired by experimental replicate. Pairwise statistical analyses are in Table 4.

**Table 4.**
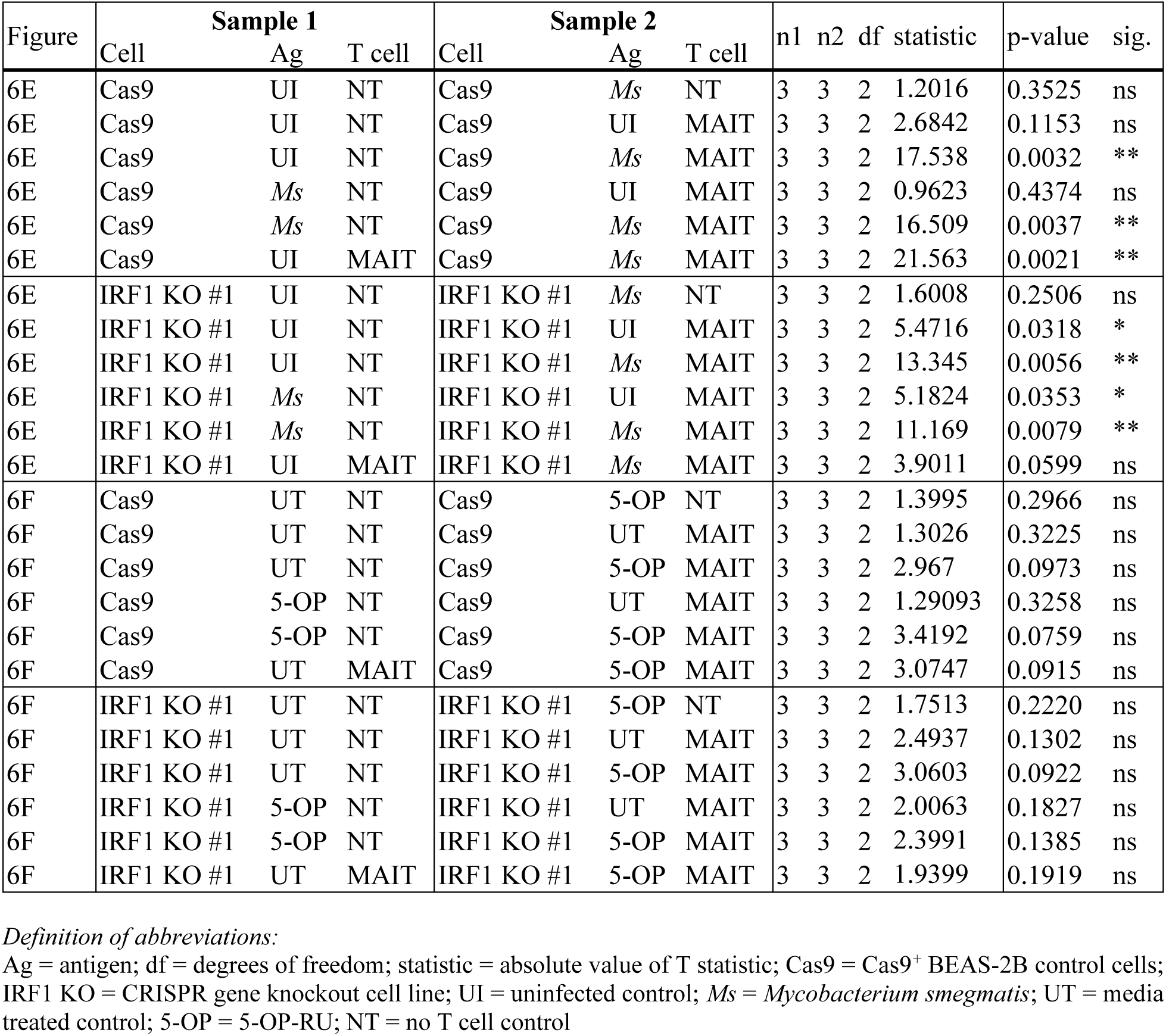
Statistics associated with Figure 6E-F.

| Figure | Sample 1 |  |  | Sample 2 |  |  | n1 | n2 | df | statistic | p-value | sig. |
| --- | --- | --- | --- | --- | --- | --- | --- | --- | --- | --- | --- | --- |
|  | Cell | Ag | T cell | Cell | Ag | T cell |  |  |  |  |  |  |
| 6E | Cas9 | UI | NT | Cas9 | *Ms* | NT | 3 | 3 | 2 | 1.2016 | 0.3525 | ns |
| 6E | Cas9 | UI | NT | Cas9 | UI | MAIT | 3 | 3 | 2 | 2.6842 | 0.1153 | ns |
| 6E | Cas9 | UI | NT | Cas9 | *Ms* | MAIT | 3 | 3 | 2 | 17.538 | 0.0032 | ** |
| 6E | Cas9 | *Ms* | NT | Cas9 | UI | MAIT | 3 | 3 | 2 | 0.9623 | 0.4374 | ns |
| 6E | Cas9 | *Ms* | NT | Cas9 | *Ms* | MAIT | 3 | 3 | 2 | 16.509 | 0.0037 | ** |
| 6E | Cas9 | UI | MAIT | Cas9 | *Ms* | MAIT | 3 | 3 | 2 | 21.563 | 0.0021 | ** |
| 6E | IRF1 KO #1 | UI | NT | IRF1 KO #1 | *Ms* | NT | 3 | 3 | 2 | 1.6008 | 0.2506 | ns |
| 6E | IRF1 KO #1 | UI | NT | IRF1 KO #1 | UI | MAIT | 3 | 3 | 2 | 5.4716 | 0.0318 | * |
| 6E | IRF1 KO #1 | UI | NT | IRF1 KO #1 | *Ms* | MAIT | 3 | 3 | 2 | 13.345 | 0.0056 | ** |
| 6E | IRF1 KO #1 | *Ms* | NT | IRF1 KO #1 | UI | MAIT | 3 | 3 | 2 | 5.1824 | 0.0353 | * |
| 6E | IRF1 KO #1 | *Ms* | NT | IRF1 KO #1 | *Ms* | MAIT | 3 | 3 | 2 | 11.169 | 0.0079 | ** |
| 6E | IRF1 KO #1 | UI | MAIT | IRF1 KO #1 | *Ms* | MAIT | 3 | 3 | 2 | 3.9011 | 0.0599 | ns |
| 6F | Cas9 | UT | NT | Cas9 | 5-OP | NT | 3 | 3 | 2 | 1.3995 | 0.2966 | ns |
| 6F | Cas9 | UT | NT | Cas9 | UT | MAIT | 3 | 3 | 2 | 1.3026 | 0.3225 | ns |
| 6F | Cas9 | UT | NT | Cas9 | 5-OP | MAIT | 3 | 3 | 2 | 2.967 | 0.0973 | ns |
| 6F | Cas9 | 5-OP | NT | Cas9 | UT | MAIT | 3 | 3 | 2 | 1.29093 | 0.3258 | ns |
| 6F | Cas9 | 5-OP | NT | Cas9 | 5-OP | MAIT | 3 | 3 | 2 | 3.4192 | 0.0759 | ns |
| 6F | Cas9 | UT | MAIT | Cas9 | 5-OP | MAIT | 3 | 3 | 2 | 3.0747 | 0.0915 | ns |
| 6F | IRF1 KO #1 | UT | NT | IRF1 KO #1 | 5-OP | NT | 3 | 3 | 2 | 1.7513 | 0.2220 | ns |
| 6F | IRF1 KO #1 | UT | NT | IRF1 KO #1 | UT | MAIT | 3 | 3 | 2 | 2.4937 | 0.1302 | ns |
| 6F | IRF1 KO #1 | UT | NT | IRF1 KO #1 | 5-OP | MAIT | 3 | 3 | 2 | 3.0603 | 0.0922 | ns |
| 6F | IRF1 KO #1 | 5-OP | NT | IRF1 KO #1 | UT | MAIT | 3 | 3 | 2 | 2.0063 | 0.1827 | ns |
| 6F | IRF1 KO #1 | 5-OP | NT | IRF1 KO #1 | 5-OP | MAIT | 3 | 3 | 2 | 2.3991 | 0.1385 | ns |
| 6F | IRF1 KO #1 | UT | MAIT | IRF1 KO #1 | 5-OP | MAIT | 3 | 3 | 2 | 1.9399 | 0.1919 | ns |

*Definition of abbreviations:*
Ag = antigen; df = degrees of freedom; statistic = absolute value of T statistic; Cas9 = Cas9<sup>+</sup> BEAS-2B control cells; IRF1 KO = CRISPR gene knockout cell line; UI = uninfected control; *Ms* = *Mycobacterium smegmatis*; UT = media treated control; 5-OP = 5-OP-RU; NT = no T cell control

Finally, we used our IRF1^−/−^ cells in this co-culture setting to demonstrate the role of IFNγ in mediating MR1 expression, antigen presentation, and MAIT cell activation. IRF1^−/−^ cells infected with *Ms* and co-cultured with MAIT cells failed to exhibit the increase in *MR1* expression seen in the Cas9^+^ control cells (Figure 6E, Table 4). We then used exogenous 5-OP-RU treatment to directly test the role of IRF1 following MR1-dependent MAIT cell activation. *MR1* transcription was enhanced in Cas9^+^ cells with 5-OP-RU and MAIT cell co-culture, confirming that TCR-stimulated MAIT cells directly led to increased *MR1* transcription (Figure 6F, Table 4). In contrast, the IRF1^−/−^ cells treated with 5-OP-RU did not express greater *MR1* transcripts in co-culture, confirming the importance of IRF1 in this pathway.

### IFNγ and IFNβ stimulate MR1 transcription by distinct mechanisms

Our data have shown thus far that IFNγ stimulates *MR1* transcription; however other inflammatory cytokines can also induce transcription. Type I interferons like IFNβ stimulate IRF1 and NLRC5 to induce MHC-Ia transcription^15–19,33,34^. Activated MAIT cells may also produce TNFα and IL-17, which have been demonstrated to stimulate transcription of inflammatory genes including *IRF1* and *HLA-A*^35–37^. To assess whether the IFNγ-induced increase in *MR1* transcription is representative of general inflammatory signaling mechanisms or specific to IFNγ stimulus, we treated BEAS-2B cells with recombinant human inflammatory cytokines IFNβ, IFNγ, IFNλ, TNFα, and IL-17. Of these cytokines, only IFNβ and IFNγ elicited a significant increase in *MR1* transcription compared to untreated controls (Figure 7A-B, Table 5).

**Figure 7.**
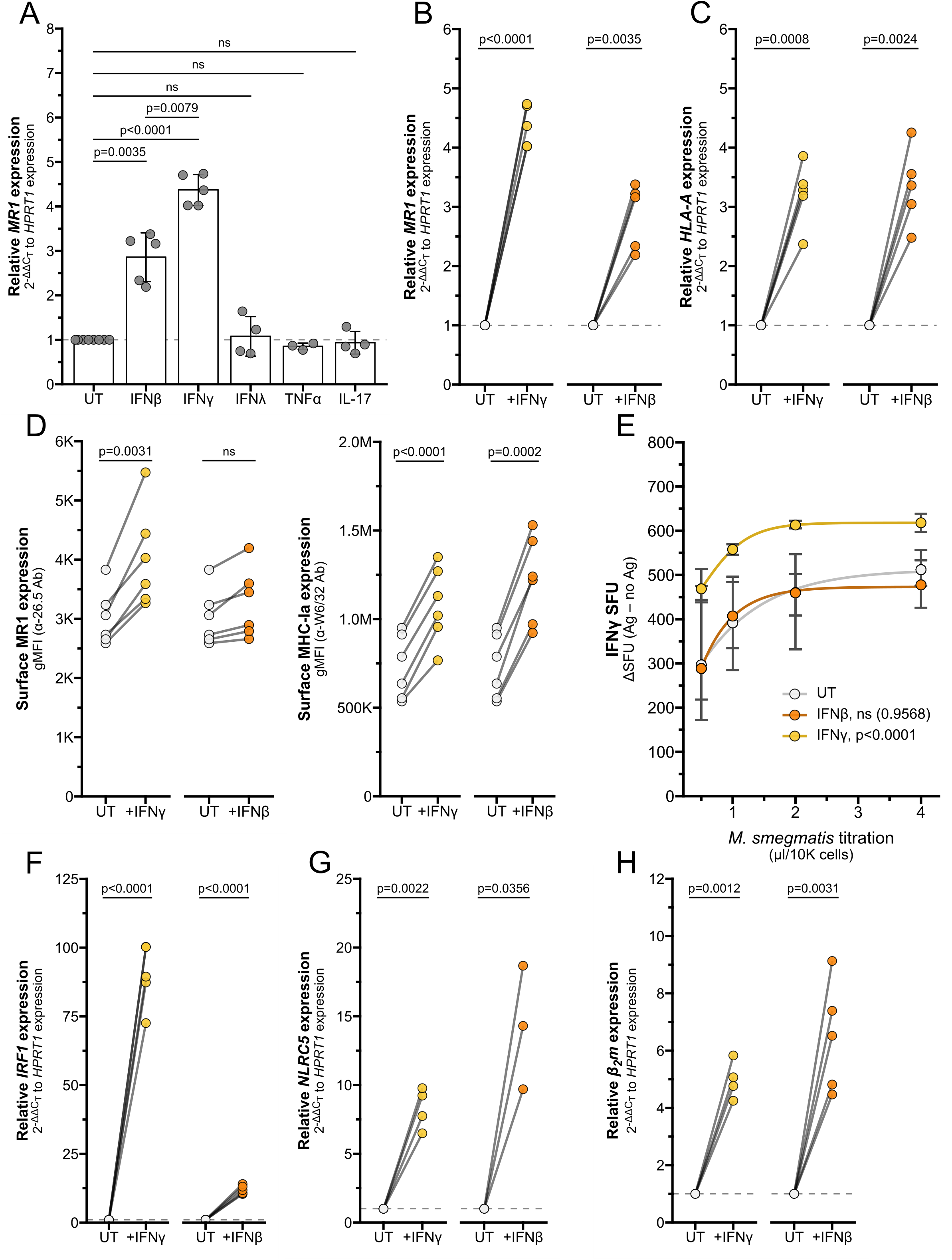
Interferons stimulate MR1 and MHC-Ia transcription through different pathways. **(A)** RT-qPCR of wildtype BEAS-2B cells treated with recombinant human cytokines for 12 hours. *MR1* expression was calculated relative to *HPRT1* expression and UT controls. **(B-C, F-H)** RT-qPCR of BEAS-2B cells treated with IFNγ or IFNβ for 12 hours. Expression of **(B)** *MR1,* **(C)** *HLA-A* **(F)** *IRF1*, **(G)** *NLRC5*, and **(H)** *β2m* were calculated relative to *HPRT1* and UT control, paired by experiment. **(D)** Flow cytometry of BEAS-2B cells treated with IFNγ or IFNβ for 12 hours. gMFI of surface MR1 (left, α-26.5 Ab) and MHC-Ia (right, α-W6/32 Ab) are paired by experimental replicate. **(E)** ELISPOT of BEAS-2B cells treated with IFNγ or IFNβ for 12 hours, infected with a titration of *M. smegmatis* for one hour, then incubated with MAIT cells overnight. Data points are average SFU of no-antigen background-subtracted IFNγ SFU. Nonlinear regression agonist response curves were computed in GraphPad Prism 10.4.0 and analyzed with extra sum-of-squares F test to compare with UT control. Statistical analyses are in Table 5.

**Table 5.**
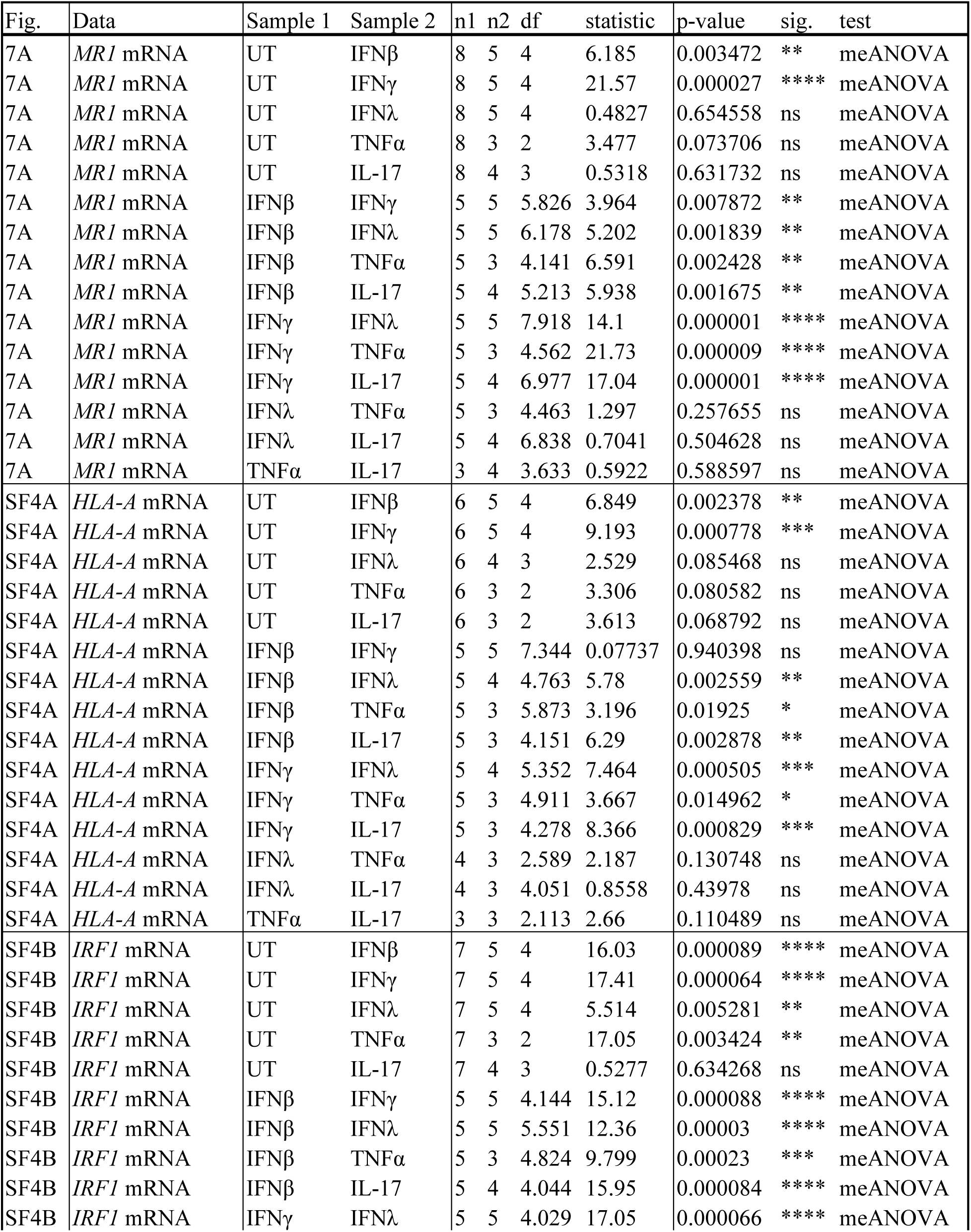
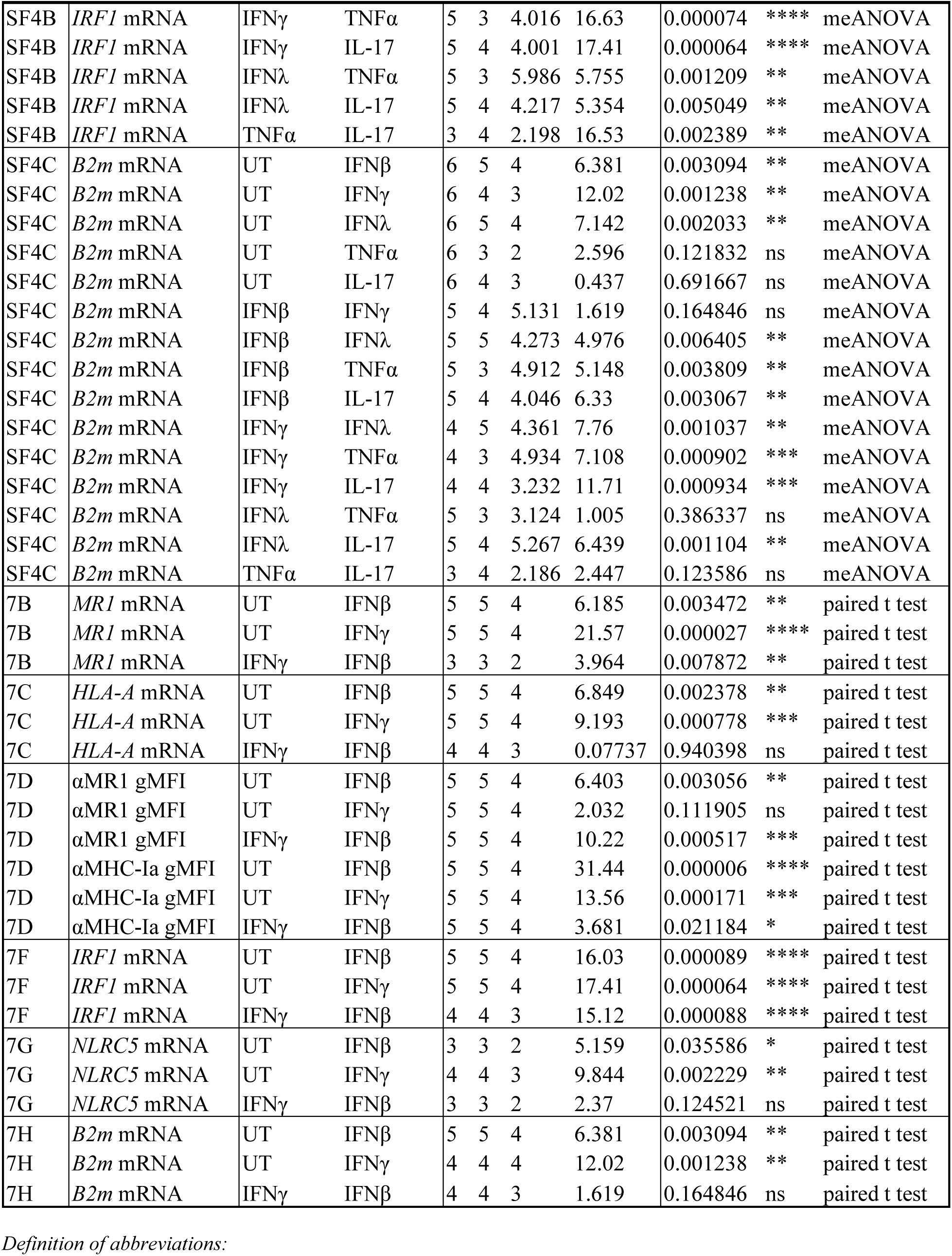
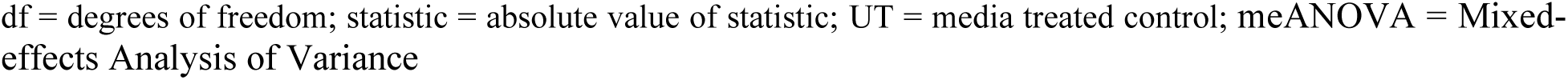
Statistics associated with Figure 7 and Supplemental Figure 4.

Interestingly, the IFNγ-mediated increase in MR1 expression was significantly greater than the increase due to IFNβ treatment, while expression of *HLA-A* and *B2m* were similarly induced by both IFNγ and IFNβ treatment (Figure 7A-C, Supplemental Figure 4, Table 5). We further investigated the role of type I and II IFNs in mediating expression of MR1 and MHC-Ia. While both IFNβ and IFNγ increased surface expression of MHC-Ia, only IFNγ led to a significant increase in surface MR1 protein expression (Figure 7D). BEAS-2B cells pre-treated with IFNγ induced significantly greater MAIT cell responses to *Ms* infection than control UT cells, while IFNβ pre-treatment did not generate a significantly different dose-response curve (Figure 7E). These data indicate that IFNγ plays the largest role in stimulating MR1 expression and function.

We quantified *IRF1* and *NLRC5* transcripts to further explore how MR1 and MHC-I expression are differentially stimulated by IFNβ and IFNγ. Although both IFNγ and IFNβ increased *IRF1* expression, the relative fold change was significantly greater with IFNγ than IFNβ (Figure 7F, Supplemental Figure 4, Table 5). Both interferons induced significant increases in *NLRC5* and *B2m* expression (Figure 7G-H, Supplemental Figure 4, Table 5). In light of our finding that IFNγ stimulates *MR1* transcription through IRF1 and not NLRC5, this magnitude of *IRF1* induction may relate to the specific induction of MR1 surface expression and antigen presentation by IFNγ and not IFNβ. Further, these results indicate that transcription of *MR1* and *HLAA* occur via distinct IFNγ-stimulated mechanisms.

Together, our data support a feed-forward model of inflammatory signaling (Figure 8). MR1 antigen presentation by an infected cell activates a MAIT cell to release IFNγ, which then acts on the airway epithelial cell to stimulate *IRF1* expression through a pSTAT1 pathway. IRF1 then binds to the *MR1* promoter to induce *MR1* transcription, leading to more MR1 protein available for antigen presentation and subsequent MAIT cell activation.

**Figure 8.**
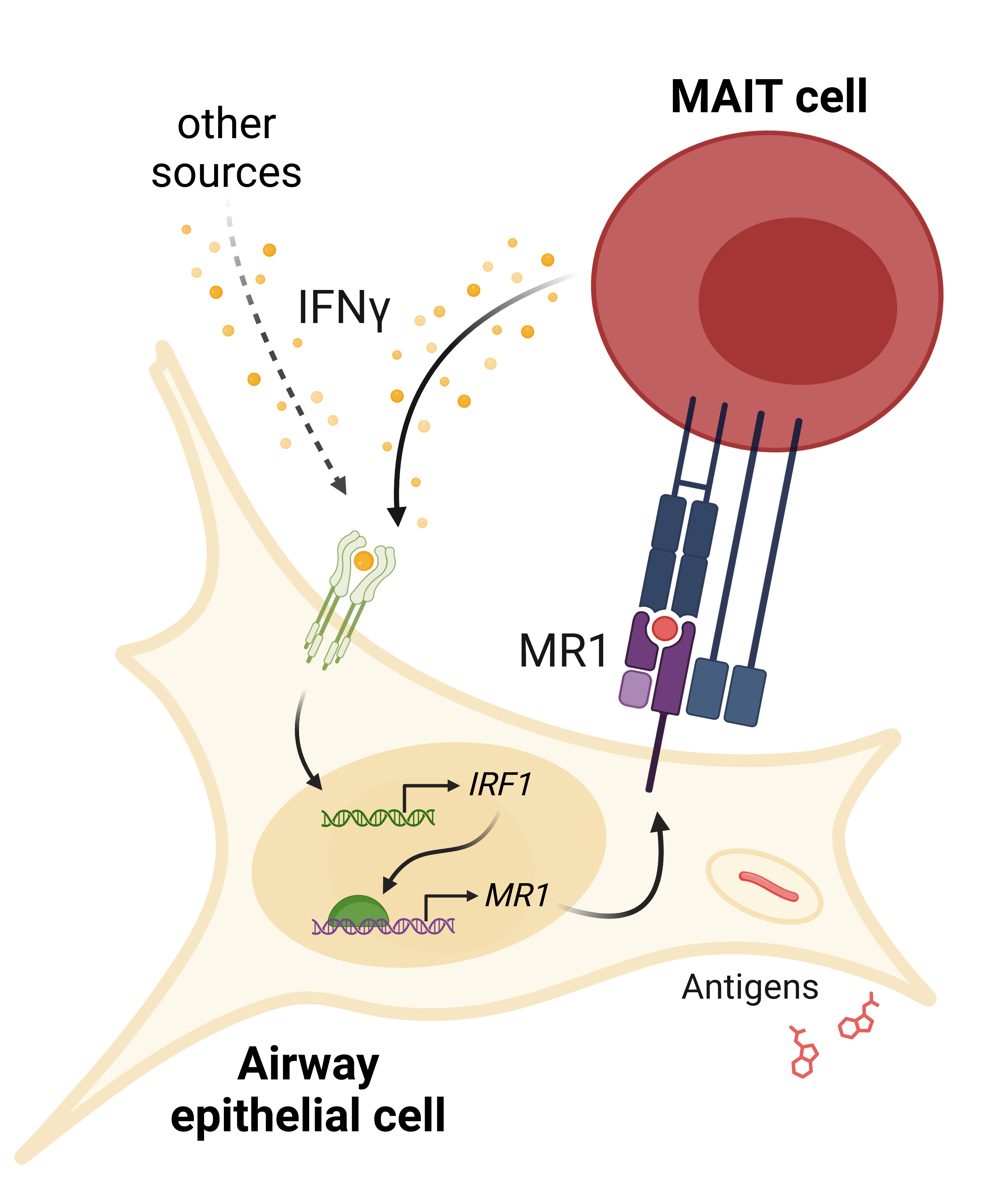
IFNγ signaling induces MR1 expression and MAIT cell activation. IFNγ from MAIT cells or other cellular sources induces *IRF1* expression and subsequent increase in *MR1* transcription. Increased MR1 expression and antigen presentation enhances MAIT cell responses to antigens from exogenous sources or pathogens like *S. pneumoniae* or *M. smegmatis*.

## Discussion

MAIT cells are key components of early infection responses. The variety of pathogens producing MR1 antigens and rapid MAIT cell effector function poise MAIT cells to bridge innate and adaptive immune responses. Strict regulation of MAIT cell activation is required to prevent inflammatory damage. Research over the past decade has defined many complementary pathways regulating MR1 intracellular localization, antigen binding, surface translocation, and protein recycling^12–14^. These studies affirm that defining the mechanisms that regulate MR1 is critical to understanding regulation of MAIT cells themselves. Only recently have we begun to appreciate the role of epigenetic regulation in controlling MR1 expression and antigen presentation. Studies of *MR1* DNA methylation and RNA expression suggest that *MR1* transcription is increased during infection^38–41^, although effector molecules from herpes simplex viruses were shown to degrade *MR1* transcripts^42–44^. Altered epigenetic regulation of *MR1* in respiratory inflammation^22–24,45^ and cancer^21,46^ illustrate the complex interplay of activation and repression signals in these diseases.

Here, we demonstrate for the first time that IFNγ stimulates *MR1* transcription. We investigated the role of two IFNγ-stimulated transcription factors, IRF1 and NLRC5, in regulating *MR1* transcription. *In-silico* analysis of the *MR1* promoter revealed potential binding sites for IRF1 and components of the NLRC5 enhanceosome^18,19^. Using both siRNA knockdown and CRISPR/Cas9 knockout systems, we observed that IFNγ-induced *MR1* transcription was dependent on IRF1, but not NLRC5. Treatment with IFNγ failed to increase MR1 surface expression and antigen presentation in IRF1^−/−^ cells. These results pointed to IRF1 as the primary driver of IFNγ-induced *MR1* transcription in our experimental conditions. The rapid increase in *IRF1* expression after IFNγ treatment is consistent with established models of IRF1 kinetics^18,47^. Recently, Rosain and colleagues performed a comprehensive characterization of two young patients with IRF1-inactivating mutations^48^. *MR1* was among the genes upregulated in primary fibroblasts from healthy controls after 8 hours of IFNγ treatment, while *MR1* was not upregulated in IFNγ-treated fibroblasts with IRF1-inactivation, STAT1-deficiency, or loss of IFNGR1/2^48^. Our co-culture experiments showed that antigen-activated MAIT cells produced sufficient IFNγ to induce *IRF1* expression in both primary AEC and BEAS-2B cells. In similar conditions with *Ms* infection or 5-OP-RU treatment and MAIT cell co-culture, IRF1^−/−^ cells failed to replicate the increase in *MR1* mRNA expression seen with control Cas9^+^ cells. We therefore concluded that MAIT cell activation acts through IRF1 to promote *MR1* transcription. The case study of IRF1-deficient patients observed only slightly lower blood MAIT cell frequencies in one individual^48^. However, both experienced persistent early childhood infections from weakly-infectious *Mycobacterium avium* and/or breakthrough infection from the BCG vaccine, consistent with impaired IFNγ immunity^48^. Although many factors may contribute to reduced immune function in these individuals, MAIT cells have been tightly linked to anti-mycobacterial immunity^49,50^. A case study of a T-bet-deficient individual with very low MAIT cells underscored the specific importance of IFNγ production by innate-like lymphocytes in controlling *Mtb* infection^51^. It is therefore possible that loss of IRF1 function could foster mycobacterial susceptibility through weaker induction of *MR1* transcription and delayed MAIT cell responses. Directly characterizing the dynamics of *MR1* expression in these individuals would shed light on this hypothesis.

We were surprised to observe that NLRC5 was not required for the IFNγ-induced increase in MR1 expression or antigen presentation, given the importance of NLRC5 in mediating *HLA* transcription^15,16,19^. Although MR1 is an MHC-I-like molecule and they share broad structural homology, the genes reside on different chromosomes and have distinct promoter features^7^. We confirmed that *HLA-A* expression was reduced in NLRC5^−/−^ cells. Since IRF1 can also induce *NLRC5* transcription, we considered whether IRF1 and NLRC5 play a synergistic role in regulating MR1 expression. While the combined loss of IRF1 and NLRC5 reduces Class I mRNA expression, we saw no further impact to MR1 expression or antigen presentation. These results prompted us to explore how MHC-Ia and MR1 transcription signaling pathways diverge. Both type I and type II IFNs are known to stimulate expression of IRF1 and MHC-Ia^15–18,33,34^. We explored whether IFNβ might induce *MR1* transcription similarly to IFNγ. Although both interferons increased *MR1* mRNA expression, IFNγ stimulated a significantly greater increase in *MR1* transcription than IFNβ. Furthermore, only IFNγ induced MR1 surface protein expression and antigen presentation to MAIT cells. Previously, Ussher *et al.* demonstrated that MAIT cell responses to fixed intact *E. coli* were significantly increased when THP1 cells were incubated with either IFNα or IFNγ overnight^52^. This type I IFN increase does not match our results. However, several other groups observed that directly stimulating MAIT cells with IFNβ or IFNα led to increased TCR-dependent and -independent MAIT cell responses to influenza virus and *Klebsiella pneumoniae* infection^53–55^. Therefore, type I interferon-induced signaling within MAIT cells may be the primary driver of the observed increase in MAIT cell responses^52^. Type III interferons signal through similar pathways as type I interferons and are critical to some mucosal inflammatory responses^56^. We failed to measure any notable increase in *MR1* or *HLA-A* expression following IFNλ treatment. We observed significant increases in *IRF1* and *β2m* expression with IFNλ; however, these increases were significantly weaker than any IFNγ- or IFNβ-induced *IRF1* or *β2m* expression. This result is consistent with research from Forero *et al.*, who found that IFNλ primarily induces tissue repair pathways and fails to stimulate *IRF1* expression^57,58^.

Others have shown that IFNγ, not IFNβ, is the primary driver of *IRF1* expression^48^. We quantified the relative increase in *IRF1* transcripts following stimulation by IFNγ or IFNβ and likewise observed the increase in *IRF1* expression was significantly higher with IFNγ compared to IFNβ. In contrast, both interferons led to similar ranges of *NLRC5* expression. One possible model of this data suggests that IFNγ signaling induces sufficient IRF1 expression to promote *MR1* transcription. IFNβ induces less *IRF1* transcription, leading to expression of NLRC5, MHC-Ia, and β₂m, but not enough to stimulate MR1 expression and function. It is tempting to speculate that the specificity of MR1 stimulation by IFNγ may help to compartmentalize immune responses to innate signaling and prevent simultaneous overstimulation of both MHC-Ia and MR1.

We also observed a slight increase in *IRF1* expression following TNFα treatment, yet no increase in *HLAA, β2m,* or *MR1* expression. It was surprising that TNFα did not stimulate any significant increase in IRF1-induced genes. The TNF receptor-associated factor 6 (TRAF6) works with cellular inhibitor of apoptosis 2 (cIAP2) to K63 ubiquitinate IRF1, leading to increased function and blocking K48 ubiquitin-mediated IRF1 proteasomal degradation^18,47,59–63^. This process, however, functions in concert with Src-family kinases following TLR4, TLR7/9, or IL-1 signaling^18,47,61–65^. It is possible that this signaling occurred in our experimental condition with primary AEC infected with TLR4 ligand-producing *S. pneumoniae*, since we observed increases in *IRF1* and *HLA-A* expression with or without MAIT cells. However, we also observed IRF1-dependent *MR1* expression following 5-OP-RU-induced MAIT cell activation, in which circumstance TLR signaling was likely inactive. Exploring the role of innate sensors and non-interferon cytokines in modulating IRF1 activity may reveal additional factors that can induce or repress *MR1* expression.

In co-culture settings, IFNγ-stimulated increases in *MR1* expression resulted in greater MR1 antigen presentation and subsequent MAIT cell activation. MAIT cells activated by MR1 antigen presentation produced sufficient IFNγ to promote *MR1* transcription and increased MR1 surface expression. This feed-forward signaling model would support the function of MAIT cells in immune surveillance and early infection response. Robust stimulation of MR1-dependent MAIT cell activation could be beneficial during infection onset, allowing minimal antigen stimulus to generate expansive and rapid proinflammatory activity. MAIT cell effector functions are well-established in priming myeloid cells and recruiting CD4^+^ and CD8^+^ T cells^66,67^. In addition to MAIT cells, a number of other cells could produce IFNγ and initiate this feed-forward loop. Local IFNγ production by professional antigen-presenting cells has been observed in infection contexts; for example, alveolar macrophages produce IFNγ during *M. tuberculosis* infection^68,69^. NK cells, ILC, airway-resident lymphocytes, and circulating lymphocytes are also known to make IFNγ in response to a variety of inflammatory stimuli as well^70,71^. Beyond cytokine signaling, toll-like receptors and C-type lectin receptors can also stimulate expression of *IRF1* and IRF1-inducible genes, suggesting that IRF1 inflammation could be mediated in response to non-interferon stimuli^72^. IFNγ and IRF1 signaling through any of these sources could then stimulate MR1 expression and activate MAIT cells, leading to enhanced inflammatory responses.

However, dysregulation of this feed-forward loop could also lead to MAIT cell-caused pathology. Inappropriate MAIT cell activation is implicated in autoimmune diseases and chronic inflammation^73,74^. Overproduction of IFNγ contributes to inflammatory lung damage and can stimulate further IFNγ production by alveolar macrophages^68,75–77^. CD8^+^ T cell infiltration is associated with increased disease severity in chronic obstructive pulmonary disease (COPD), and IFNγ signaling is increased in the lungs of COPD patients^77–80^. We previously characterized expression of MR1 and activation of MAIT cells in the context of COPD and cigarette smoke^22^. We found that primary AEC from COPD donors induced greater MAIT cell activation in the absence of antigen than healthy AEC. *MR1* transcriptional expression was increased in AEC^22^ and PBMC^24^ from infected COPD donors. It is possible that the increased IFNγ present in the COPD airway environment could stimulate *MR1* expression and lead to increased MAIT cell activity via the proposed feed-forward signaling loop. IFNγ signaling in response to inappropriate stimuli (e.g. antigens from commensal microbes) could also induce *MR1* transcription and promote ligand-driven MAIT cell inflammatory pathology.

Given the potential for inflammatory damage due to this feed-forward loop, we hypothesize that a dampening mechanism exists to turn off this pathway. Constantin *et al.* revealed a potential role for ERK1/2 kinases in suppressing *MR1* expression in melanoma, indicating repression mechanisms can modulate *MR1* transcription^46^. Specifically, they found the transcription factor ELF1 binds to the *MR1* promoter to stimulate *MR1* expression. ERK1/2, members of the MAPK/MEK signaling cascade, inhibited ELF1 function and subsequent *MR1* transcription. The authors suggested ELF1 inhibition may occur through post-translational modification performed by a downstream intermediary protein^46^. Signaling through MEK/ERK was recently demonstrated to inhibit IRF1 expression and activity in TLR-stimulated macrophages; however, ELF1 mediates antiviral activity in airway epithelial cells independent of interferon and IRF1 transcriptional activity^81,82^. Multiple distinct mechanisms of *MR1* transcriptional activation could serve several functions: flexible induction of *MR1* expression and function in the context of distinct stimuli, complementary activation to rapidly enhance MAIT cell responses, and/or as a checkpoint requiring a secondary signal to prevent overactivation. It is well-documented that pathogens target IFNγ signaling and MHC transcription mechanisms to evade immune recognition^34,83,84^. Several MHC-Ia post-transcriptional repression mechanisms have been identified, including through IRF1 degradation or downregulation of NLRC5 expression^16,85,86^. In human fibroblasts, *MR1* transcripts were degraded by an RNase protein from herpes simplex virus types 1 and 2, although this mechanism was not specific to *MR1*^42,43^. A greater understanding of how *MR1* transcription is regulated could shed light on these host-pathogen dynamics.

Put together, this work demonstrates that IFNγ signaling stimulates *MR1* transcription, surface expression, and antigen presentation. Understanding the mechanisms of *MR1* transcriptional regulation may provide insights into broader immune signaling networks and better inform our knowledge of the roles MR1 and MAIT cells play in infection and inflammatory diseases.

## Materials & Methods

### Human subjects

This study was conducted according to the principles expressed in the Declaration of Helsinki. Study participants, protocols and consent forms were approved by Oregon Health & Science University Institutional Review Board (IRB00000186). Written and informed consent was obtained from all donors. Human participants are not directly involved in the study. Healthy adults were recruited from among employees at Oregon Health & Science University as previously described to obtain human serum^87^.

### Cells and bacteria

Primary airway epithelial cells (AEC) were purchased from Lonza Biosciences or harvested from deceased human donor lung tissue through the Cascade Alliance (formerly Pacific Northwest Transplant Bank) as previously described^22,88^. The healthy donor AEC from ^22^ were likewise grown in Bronchial Epithelial Growth Media (“BEGM”, CC-3170) and harvested using ReagentPack Subculture reagents (CC-5034) per manufacturer’s protocols (Lonza).

The BEAS-2B bronchial epithelial cell line (CRL-9609, American Type Culture Collection) was grown in DMEM medium (Gibco) supplemented with L-glutamine (25030164, Life Technologies) and 10% heat-inactivated fetal bovine serum (“DMEM-FBS”). BEAS-2B cells overexpressing MR1-GFP under a tetracycline-inducible promoter (“BEAS-2B:doxMR1-GFP”)^11^ were similarly cultured in DMEM-FBS. Expression of MR1-GFP was induced with doxycycline for 16 hours prior to harvest. BEAS-2B cells stably expressing Cas9^89^ were grown in DMEM-FBS and used to generate CRISPR knockouts.

The MR1-restricted T cell clone (D426G11) was generated and expanded in RPMI medium (Gibco) supplemented with L-glutamine and 10% heat-inactivated human serum (“RPMI-HuS”) as previously described^2,87^.

*Streptococcus pneumoniae*^90^ and *Mycobacterium smegmatis* Mc^2^155 (ATCC) were grown as described in the supplement of ^22^ and used from frozen stocks. At late log phase, *M. smegmatis* were pelleted and the supernatant was passed through a syringe-driven 0.22 μm filter and frozen for use as antigen in ELISPOT assays.

### Generation of stable CRISPR/Cas9 IRF1 or NLRC5 knockout BEAS-2B cells

We generated IRF1^−/−^ and NLRC5^−/−^ CRISPR knockout BEAS-2B cells as previously described^89^. Early passage Cas9^+^ BEAS-2B cells were transduced with sgRNA constructs targeting IRF1 (CRISPR845545_LV, ThermoFisher) or NLRC5 (CRISPR1120312_LV, ThermoFisher) in the presence of 200 μg Polybrene (Sigma). Following puromycin selection, monoclonal populations were produced by limiting dilution and screened by Western blot or ELISPOT. We validated genomic editing by Sanger sequencing. DNA was isolated from control Cas9^+^, IRF1^−/−^, and NLRC5^−/−^ BEAS-2B clones using the QIAamp DNA Micro Kit (Qiagen) and amplified by PCR. The OHSU Vollum Institute DNA Sequencing Core performed Sanger sequencing and the resulting sequences were analyzed by TIDE^91^ and ICE^92^.

### Reagents and antibodies

6-formylpterin (6-FP, Schirck’s Laboratories) was suspended in 0.01 M NaOH and used at a final concentration of 100 μM. 5-(2-oxopropylideneamino)-6-d-ribitylaminouracil (5-OP-RU) was freshly prepared from equal volumes of 32 mM 5-amino-6-d-ribitylaminouracil (5-A-RU)*HCl (OHSU Medicinal Chemistry Core)^93^ and 650 mM methylglyoxal (Sigma) exactly following the second method described in ^89^ and used at a final concentration of 500 pM. Phytohemagglutinin PHA-L (L4144 Sigma) was suspended in RPMI-HuS and used at 1 μg/well. Doxycycline (Sigma) was suspended in sterile water and used at 2 μg/ml.

Recombinant human cytokines were reconstituted in sterile water and supplemented with bovine serum albumin as per manufacturer recommendations. Final concentrations used were: 66 ng/ml IFNγ (R&D Systems 285-IF-100), 66 ng/ml IFNβ (R&D Systems 8199-IF-010), 132 ng/ml IFNλ (PeproTech 300-02K), 66 ng/ml TNFα (R&D Systems 10291-TA-050), and 66 ng/ml IL-17 (PeproTech 200-17). Cells were treated with cytokines for 12 hours unless otherwise noted.

Antibodies used for ELISPOT assays: α-IFNγ (1-D1K, Mabtech) and alkaline phosphatase-conjugated secondary antibody (7-B6-1-ALP, Mabtech). Antibodies used for Western blot: α-IRF1 (D5E4, Cell Signaling Technology), α-Vinculin (V284, Bio-Rad). Antibodies used for flow cytometry: α-MR1 (26.5, conjugated to APC, Biolegend), α-HLA-A,B,C (W6/32, conjugated to APC, Biolegend), IgG2a isotype (MOPC-173, conjugated to APC, Biolegend), α-phospho-STAT1 (KIKSI0803, conjugated to PE, eBioscience).

### Co-culture experiments

Primary AEC were infected with *S. pneumoniae* (20 MOI) in antibiotic-free BEGM. After 1 hour, AEC were washed with PBS to remove non-adhered bacteria, then MAIT cells were added at a 1:1 ratio in BEGM complete with gentamycin-amphotericin (GA-1000, Lonza). BEAS-2B cells in antibiotic-free DMEM-FBS were infected with *M. smegmatis* or treated with 6-FP or 5-OP-RU for 1 hour, washed with PBS, then MAIT cells were added at a 1:1 ratio in DMEM-FBS with gentamycin. Following overnight co-culture, wells were extensively washed with PBS to remove MAIT cells before harvesting AEC or BEAS-2B cells.

### Real-time quantitative PCR (RT-qPCR)

Cell pellets washed with PBS were either used fresh or stored dry at −80°C before thawing in 37°C water bath. RNA was isolated using the RNEasy Plus kit (Qiagen) and cDNA was synthesized using the High Capacity cDNA Reverse Transcription Kit (Life Technologies) as per the manufacturers’ protocols. RT-qPCR was performed using TaqMan (Applied Biosystems) gene expression assays: *HPRT1* (Hs02800695_m1), *MR1* (Hs01042278_m1), *HLA-A,H* (Hs01058806_g1), *IRF1* (Hs00971965_m1), *NLRC5* (Hs01072123_m1), and *β2m* (Hs00187842_m1). Gene expression data were normalized to internal control *HPRT1* and relative expression levels for each target gene were determined using the 2^−ΔΔCt^ method^94^. Some uninfected AEC *HPRT1* and *MR1* data were used as controls in ^22^.

### Flow cytometry

To quantify surface expression of MR1 and MHC-I, AEC and BEAS-2B cells were treated as indicated and harvested. Samples were blocked in FACS buffer containing 2% heat-inactivated human serum, 2% heat-inactivated goat serum, and 0.5% heat-inactivated FBS for 30 minutes on ice, then stained with APC-conjugated IgG2a, α-MR1, or α-HLA-A,B,C antibody for 40 minutes. For pSTAT1 staining, cells were permeabilized with 0.2% saponin during the blocking step. Cells were washed with PBS and fixed with 1% paraformaldehyde, then analyzed with a Beckman Coulter CytoflexS. All analyses were performed using FlowJo10 (TreeStar).

### Enzyme-linked immunospot (ELISPOT) assays

IFNγ ELISPOT assays were performed as previously described^95^ with the following modifications: ELISPOT plates (MSHAS4510, MilliporeSigma) were coated overnight with α-IFNγ antibody, then washed and blocked for 1 hour in RPMI-HuS. BEAS-2B cells were seeded in duplicate (1×10^5^ cells/well) and infected with *M. smegmatis*, treated with a titration of *M. smegmatis* supernatant, or incubated with control PHA or RPMI-HuS medium for 1 hour at 37°C. D426G11 MAIT cell clones were added at a 1:1 ratio in RPMI-HuS with gentamycin for overnight incubation at 37°C. Following extensive washing with PBS-0.05% Tween 20, plates were incubated with ALP secondary antibody for 2 hours before additional washing and colorimetric development. IFNγ spot-forming units (SFU) were quantified by AID ELISPOT reader. For experiments with cytokine pre-treatment, BEAS-2B cells were seeded in 6-well plates and treated with cytokines for 12 hours, then washed 3 times with PBS to remove any excess cytokine before harvesting and seeding into ELISPOT plate.

### siRNA gene silencing

Gene silencing in wildtype, Cas9^+^, or NLRC5^−/−^ BEAS-2B cells was performed through nucleofection as in ^96^ and following the Amaxa Cell Line Nucleofector Kit T (Lonza) protocols. In brief, 2 μg total of Missense (4390843, ThermoFisher), IRF1 (s7501, ThermoFisher), and/or NLRC5 (s38591, ThermoFisher) siRNA were added to 1×10^6^ cells and transfected by the Amaxa Nucleofector 2b machine (Lonza) using program G-016. Cells were incubated for 48 hour before use in assays. Efficiency of gene silencing was validated by RT-qPCR.

### Transcription factor binding sites

Putative transcription factor binding sites were acquired through the Eukaryotic Promoter Database browser using the Search Motif Tool to perform on-the-fly scanning for transcription factor motifs using the FindM tool from the Signal Search Analysis (SSA) Server toolkit^28,97–99^.

### Data analysis

All data were analyzed using Prism (GraphPad) and plots were generated using R 4.4.0 and packages such as tidyverse, ggprism, and rstatix. Statistical significance was determined as indicated by two-tailed unpaired or pairwise t tests, using α=0.05.

## Acknowledgements

We thank the staff at Cascade Alliance (formerly Pacific Northwest Transplant Bank) for procuring the human lung and airway tissue samples used in this study. The D426G11 MAIT cell clone was a kind gift from Dr. David Lewinsohn. We thank the OHSU Vollum Institute DNA Sequencing Core and the OHSU Flow Cytometry Core for their support. We also thank Dr. Laurisa Ankley and Dr. Andew Olive for their technical advice along with Savannah McBride and Dr. Fikadu Tafesse for their aid in generating Cas9^+^ BEAS-2B cells. We are grateful to Dr. Georgiana Purdy, Dr. Elly Karamooz, and Dr. Corinna Kulicke for their scientific guidance.

## Supplemental Figure Legends

**Supplemental Figure 1. Associated with Figure 1**.

**(A)** RT-qPCR of RNA isolated from primary human AECs (n=5) infected with *S. pneumoniae* (*Sp*) for one hour and incubated overnight with MAIT cell clone. *MR1* expression was calculated relative to *HPRT1* expression and uninfected no-MAIT (UI-) controls, paired by individual donor.

MR1 **(B)** mRNA and **(C)** surface expression of BEAS-2B cells infected with *M. smegmatis* (*Ms*) for one hour and incubated overnight with MAIT cell clone. **(B)** RT-qPCR of *MR1* expression was calculated relative to *HPRT1* expression and UI-control, paired by experimental replicate. **(C)** gMFI of surface MR1 stained with α-MR1 26.5 Ab, paired by experimental replicate.

MR1 **(D)** mRNA and **(E)** surface expression of BEAS-2B cells treated with 5-OP-RU (left, “5-OP”) or 6-FP (right) for one hour and incubated overnight with MAIT cell clone. **(D)** RT-qPCR of *MR1* expression was calculated relative to *HPRT1* expression and media no-MAIT (UT-) control, paired by experimental replicate. **(E)** gMFI of surface MR1 stained with α-26.5 Ab, paired by experimental replicate.

Pairwise statistical analyses are in Table 1.

**Supplemental Figure 2. Associated with Figure 4**.

**(A-B)** RT-qPCR of BEAS-2B cells treated with IRF1, NLRC5, and/or missense siRNA as indicated for 36 hours, then incubated with IFNγ for 12 hours. **(A)** *IRF1* and **(B)** *NLRC5* expression were calculated relative to *HPRT1* expression and missense UT control, paired by experimental replicate.

**(C-D)** RT-qPCR of Cas9^+^ or NLRC5^−/−^ BEAS-2B cells treated with IRF1 or missense siRNA for 36 hours, then incubated with IFNγ for 12 hours. Gene expression of **(C)** NLRC5^−/−^ clone #1 or **(D)** NLRC5^−/−^ clone #2 were calculated relative to *HPRT1* expression and Cas9^+^ or NLRC5^−/−^ clone missense UT controls, paired by experimental replicate.

**(E)** Flow cytometry of cells from (C-D). gMFI of surface MR1 (α-26.5 Ab) are from IFNγ-treated Cas9^+^ (left), NLRC5^−/−^ clone #1 (middle), and NLRC5^−/−^ clone #2 (right).

**(F)** RT-qPCR of Cas9^+^ or IRF1^−/−^ clone #1 BEAS-2B cells treated with IFNγ for 12 hours. *MR1* expression was calculated relative to *HPRT1* expression and Cas9^+^ or IRF1^−/−^ clone #1 UT controls, paired by experimental replicate.

**(G)** Flow cytometry of Cas9^+^ or IRF1^−/−^ clone #2 BEAS-2B cells treated with IFNγ for 12 hours. gMFI of surface MR1 (left, α-26.5 Ab) and MHC-Ia (left, α-W6/32 Ab) are paired by experimental replicate.

Statistical analyses are in Table 2.

**Supplemental Figure 3. Associated with Figure 5 and Figure 6**.

**(A)** Flow cytometry of primary human AECs infected with *S. pneumoniae* (*Sp*) for one hour and incubated overnight with MAIT cell clone. gMFI of stained pSTAT1 expression is paired by individual donor (n=3).

**(B-D, G)** RT-qPCR of primary human AECs infected with *S. pneumoniae* (*Sp*) for one hour and incubated overnight with MAIT cell clone. Expression of **(B)** *β2m,* **(C)** *NLRC5* **(D)** *HLA-A*, and

**(G)** *IRF1* were calculated relative to *HPRT1* expression and UI-control, paired by individual donor (n=4 (B) or n=5 (C, D, G) donors).

**(E-F, H-I)** RT-qPCR of wildtype BEAS-2B cells treated as indicated below. Gene expression was calculated relative to *HPRT1* expression and UI- or UT-controls, paired by experiment.

**(E)** *HLA-A* and **(H)** *IRF1* expression of BEAS-2B cells infected with *M. smegmatis* (*Ms*) for one hour and incubated overnight with MAIT cell clone.

**(F)** *HLA-A* and **(I)** *IRF1* expression of BEAS-2B cells treated with 5-OP-RU (left) or 6-FP (right) for one hour and incubated overnight with MAIT cell clone. Pairwise statistical analyses are in Table 3.

**Supplemental Figure 4. Associated with Figure 7**.

**(A-C)** RT-qPCR of wildtype BEAS-2B cells treated with recombinant human cytokines for 12 hours. Gene expression was calculated relative to *HPRT1* expression and UT controls.

Statistical analyses are in Table 5.

## References

1. Treiner, E., Duban, L., Bahram, S., Radosavljevic, M., Wanner, V., Tilloy, F., Affaticati, P., Gilfillan, S. & Lantz, O. Selection of evolutionarily conserved mucosal-associated invariant T cells by MR1. Nature 422, 164–9 (2003).

2. Gold, M. C., Cerri, S., Smyk-Pearson, S., Cansler, M. E., Vogt, T. M., Delepine, J., Winata, E., Swarbrick, G. M., Chua, W. J., Yu, Y. Y., Lantz, O., Cook, M. S., Null, M. D., Jacoby, D. B., Harriff, M. J., Lewinsohn, D. A., Hansen, T. H. & Lewinsohn, D. M. Human mucosal associated invariant T cells detect bacterially infected cells. PLoS Biol. 8, e1000407 (2010).

3. Le Bourhis, L., Martin, E., Peguillet, I., Guihot, A., Froux, N., Core, M., Levy, E., Dusseaux, M., Meyssonnier, V., Premel, V., Ngo, C., Riteau, B., Duban, L., Robert, D., Huang, S., Rottman, M., Soudais, C. & Lantz, O. Antimicrobial activity of mucosal-associated invariant T cells. Nat. Immunol. 11, 701–8 (2010).

4. Meermeier, E. W., Harriff, M. J., Karamooz, E. & Lewinsohn, D. M. MAIT cells and microbial immunity. Immunol Cell Biol 96, 607–617 (2018).

5. Hinks, T. S. C. & Zhang, X.-W. MAIT Cell Activation and Functions. Front. Immunol. 0, (2020).

6. Kjer-Nielsen, L., Patel, O., Corbett, A. J., Le Nours, J., Meehan, B., Liu, L., Bhati, M., Chen, Z., Kostenko, L., Reantragoon, R., Williamson, N. A., Purcell, A. W., Dudek, N. L., McConville, M. J., O’Hair, R. A., Khairallah, G. N., Godfrey, D. I., Fairlie, D. P., Rossjohn, J. & McCluskey, J. MR1 presents microbial vitamin B metabolites to MAIT cells. Nature 491, 717–23 (2012).

7. Riegert, P., Wanner, V. & Bahram, S. Genomics, isoforms, expression, and phylogeny of the MHC class I-related MR1 gene. J. Immunol. 161, 4066–4077 (1998).

8. Harriff, M. J., Karamooz, E., Burr, A., Grant, W. F., Canfield, E. T., Sorensen, M. L., Moita, L. F. & Lewinsohn, D. M. Endosomal MR1 Trafficking Plays a Key Role in Presentation of *Mycobacterium tuberculosis* Ligands to MAIT Cells. PLoS Pathog 12, e1005524 (2016).

9. McWilliam, H. E., Eckle, S. B., Theodossis, A., Liu, L., Chen, Z., Wubben, J. M., Fairlie, D. P., Strugnell, R. A., Mintern, J. D., McCluskey, J., Rossjohn, J. & Villadangos, J. A. The intracellular pathway for the presentation of vitamin B-related antigens by the antigen-presenting molecule MR1. Nat. Immunol. 17, 531–7 (2016).

10. Karamooz, E., Harriff, M. J., Narayanan, G. A., Worley, A. H. & Lewinsohn, D. M. MR1 recycling and blockade of endosomal trafficking reveal distinguishable antigen presentation pathways between *Mycobacterium tuberculosis* infection and exogenously delivered antigens. Sci. Rep. 9, 4797 (2019).

11. Huber, M. E., Kurapova, R., Heisler, C. M., Karamooz, E., Tafesse, F. G. & Harriff, M. J. Rab6 regulates recycling and retrograde trafficking of MR1 molecules. Sci. Rep. 10, (2020).

12. Kulicke, C., Karamooz, E., Lewinsohn, D. & Harriff, M. Covering All the Bases: Complementary MR1 Antigen Presentation Pathways Sample Diverse Antigens and Intracellular Compartments. Front Immunol 11, 2034 (2020).

13. Lamichhane, R. & Ussher, J. E. Expression and trafficking of MR1. Immunology 151, 270–279 (2017).

14. McWilliam, H. E. G. & Villadangos, J. A. How MR1 Presents a Pathogen Metabolic Signature to Mucosal-Associated Invariant T (MAIT) Cells. Trends Immunol. 38, 679–689 (2017).

15. van dan Elsen, P. J. Expression Regulation of Major Histocompatibility Complex Class I and Class II Encoding Genes. Front. Immunol. 2, (2011).

16. Jongsma, M. L. M., Guarda, G. & Spaapen, R. M. The regulatory network behind MHC class I expression. Mol. Immunol. 113, 16–21 (2019).

17. Kalvakolanu, D. V., Nallar, S. C. & Kalakonda, S. Interferons: Cellular and Molecular Biology of Their Actions. in Encyclopedia of Cancer (Third Edition) (eds. Boffetta, P. & Hainaut, P.) 286–312 (Academic Press, 2019). doi:10.1016/B978-0-12-801238-3.96116-6.

18. Wang, L., Zhu, Y., Zhang, N., Xian, Y., Tang, Y., Ye, J., Reza, F., He, G., Wen, X. & Jiang, X. The multiple roles of interferon regulatory factor family in health and disease. Sig Transduct Target Ther 9, 1–48 (2024).

19. Cho, S. X., Vijayan, S., Yoo, J.-S., Watanabe, T., Ouda, R., An, N. & Kobayashi, K. S. MHC class I transactivator NLRC5 in host immunity, cancer and beyond. Immunology 162, 252–261 (2021).

20. Seshadri, C., Thuong, N. T., Mai, N. T., Bang, N. D., Chau, T. T., Lewinsohn, D. M., Thwaites, G. E., Dunstan, S. J. & Hawn, T. R. A polymorphism in human MR1 is associated with mRNA expression and susceptibility to tuberculosis. Genes Immun 18, 8–14 (2017).

21. Kubica, P., Lara-Velazquez, M., Bam, M., Siraj, S., Ong, I., Liu, P., Priya, R., Salamat, S., Brutkiewicz, R. R. & Dey, M. MR1 overexpression correlates with poor clinical prognosis in glioma patients. Neuro-Oncology Advances 3, vdab034 (2021).

22. Huber, M. E., Larson, E., Lust, T. N., Heisler, C. M. & Harriff, M. J. Chronic Obstructive Pulmonary Disease and Cigarette Smoke Lead to Dysregulated Mucosal-associated Invariant T-Cell Activation. Am J Respir Cell Mol Biol 68, 90–102 (2023).

23. Sundar, I. K., Yin, Q., Baier, B. S., Yan, L., Mazur, W., Li, D., Susiarjo, M. & Rahman, I. DNA methylation profiling in peripheral lung tissues of smokers and patients with COPD. Clin Epigenetics 9, 38 (2017).

24. Szabo, M., Sarosi, V., Baliko, Z., Bodo, K., Farkas, N., Berki, T. & Engelmann, P. Deficiency of innate-like T lymphocytes in chronic obstructive pulmonary disease. Respir. Res. 18, 197 (2017).

25. McWilliam, H. E., Birkinshaw, R. W., Villadangos, J. A., McCluskey, J. & Rossjohn, J. MR1 presentation of vitamin B-based metabolite ligands. Curr. Opin. Immunol. 34, 28–34 (2015).

26. Dreos, R., Ambrosini, G., Périer, R. C. & Bucher, P. The Eukaryotic Promoter Database: expansion of EPDnew and new promoter analysis tools. Nucleic Acids Research 43, D92–D96 (2015).

27. Meylan, P., Dreos, R., Ambrosini, G., Groux, R. & Bucher, P. EPD in 2020: enhanced data visualization and extension to ncRNA promoters. Nucleic Acids Res 48, D65–D69 (2020).

28. Rauluseviciute, I., Riudavets-Puig, R., Blanc-Mathieu, R., Castro-Mondragon, J. A., Ferenc, K., Kumar, V., Lemma, R. B., Lucas, J., Chèneby, J., Baranasic, D., Khan, A., Fornes, O., Gundersen, S., Johansen, M., Hovig, E., Lenhard, B., Sandelin, A., Wasserman, W. W., Parcy, F. & Mathelier, A. JASPAR 2024: 20th anniversary of the open-access database of transcription factor binding profiles. Nucleic Acids Research 52, D174–D182 (2024).

29. Sachini, N. & Papamatheakis, J. NF-Y and the immune response: Dissecting the complex regulation of *MHC* genes. Biochimica et Biophysica Acta (BBA) - Gene Regulatory Mechanisms 1860, 537–542 (2017).

30. Ning, S., Pagano, J. S. & Barber, G. N. IRF7: activation, regulation, modification and function. Genes Immun 12, 399–414 (2011).

31. Sari, G., Dhatchinamoorthy, K., Orellano-Ariza, L., Ferreira, L. M., Brehm, M. A. & Rock, K. IRF2 loss is associated with reduced MHC I pathway transcripts in subsets of most human cancers and causes resistance to checkpoint immunotherapy in human and mouse melanomas. Journal of Experimental & Clinical Cancer Research : CR 43, 276 (2024).

32. Barrat, F. J., Crow, M. K. & Ivashkiv, L. B. Interferon target-gene expression and epigenomic signatures in health and disease. Nat Immunol 20, 1574–1583 (2019).

33. McNab, F., Mayer-Barber, K., Sher, A., Wack, A. & O’Garra, A. Type I interferons in infectious disease. Nat Rev Immunol 15, 87–103 (2015).

34. Lee, A. J. & Ashkar, A. A. The Dual Nature of Type I and Type II Interferons. Frontiers in Immunology 9, 2061 (2018).

35. Han, L.-T., Hu, J.-Q., Ma, B., Wen, D., Zhang, T.-T., Lu, Z.-W., Wei, W.-J., Wang, Y.-L., Wang, Y., Liao, T. & Ji, Q.-H. IL-17A increases MHC class I expression and promotes T cell activation in papillary thyroid cancer patients with coexistent Hashimoto’s thyroiditis. Diagnostic Pathology 14, 52 (2019).

36. Bonelli, M., Dalwigk, K., Platzer, A., Olmos Calvo, I., Hayer, S., Niederreiter, B., Holinka, J., Sevelda, F., Pap, T., Steiner, G., Superti-Furga, G., Smolen, J. S., Kiener, H. P. & Karonitsch, T. IRF1 is critical for the TNF-driven interferon response in rheumatoid fibroblast-like synoviocytes. Exp Mol Med 51, 1–11 (2019).

37. Liu, Z. Molecular mechanism of TNF signaling and beyond. Cell Res 15, 24–27 (2005).

38. Looney, M., Lorenc, R., Halushka, M. K. & Karakousis, P. C. Key macrophage responses to infection with *Mycobacterium tuberculosis* are co-regulated by microRNAs and DNA methylation. Front. Immunol. 12, (2021).

39. Pacis, A., Tailleux, L., Morin, A. M., Lambourne, J., MacIsaac, J. L., Yotova, V., Dumaine, A., Danckaert, A., Luca, F., Grenier, J.-C., Hansen, K. D., Gicquel, B., Yu, M., Pai, A., He, C., Tung, J., Pastinen, T., Kobor, M. S., Pique-Regi, R., Gilad, Y. & Barreiro, L. B. Bacterial infection remodels the DNA methylation landscape of human dendritic cells. Genome Res. 25, 1801–1811 (2015).

40. Cizmeci, D., Dempster, E. L., Champion, O. L., Wagley, S., Akman, O. E., Prior, J. L., Soyer, O. S., Mill, J. & Titball, R. W. Mapping epigenetic changes to the host cell genome induced by Burkholderia pseudomallei reveals pathogen-specific and pathogen-generic signatures of infection. Sci. Rep. 6, 30861 (2016).

41. Liu, Y., Zhu, P., Wang, W., Tan, X., Liu, C., Chen, Y., Pei, R., Cheng, X., Wu, M., Guo, Q., Liang, H., Liang, Z., Liu, J., Xu, Y., Wu, X. & Weng, X. Mucosal-Associated Invariant T Cell Dysregulation Correlates With Conjugated Bilirubin Level in Chronic HBV Infection. Hepatology 73, 1671 (2021).

42. McSharry, B. P., Samer, C., McWilliam, H. E. G., Ashley, C. L., Yee, M. B., Steain, M., Liu, L., Fairlie, D. P., Kinchington, P. R., McCluskey, J., Abendroth, A., Villadangos, J. A., Rossjohn, J. & Slobedman, B. Virus-Mediated Suppression of the Antigen Presentation Molecule MR1. Cell reports 30, 2948 (2020).

43. Samer, C., McWilliam, H. E. G., McSharry, B. P., Velusamy, T., Burchfield, J. G., Stanton, R. J., Tscharke, D. C., Rossjohn, J., Villadangos, J. A., Abendroth, A. & Slobedman, B. Multi-targeted loss of the antigen presentation molecule MR1 during HSV-1 and HSV-2 infection. iScience 27, 108801 (2024).

44. Tormanen, K., Matundan, H. H., Wang, S., Jaggi, U., Mott, K. R. & Ghiasi, H. Small Noncoding RNA (sncRNA1) within the Latency-Associated Transcript Modulates Herpes Simplex Virus 1 Virulence and the Host Immune Response during Acute but Not Latent Infection. Journal of Virology 96, e00054–22 (2022).

45. Rebuli, M. E., Glista-Baker, E., Hoffman, J. R., Duffney, P. F., Robinette, C., Speen, A. M., Pawlak, E. A., Dhingra, R., Noah, T. L. & Jaspers, I. Electronic-Cigarette Use Alters Nasal Mucosal Immune Response to Live-attenuated Influenza Virus. A Clinical Trial. Am. J. Respir. Cell Mol. Biol. 64, 126–137 (2021).

46. Constantin, D., Nosi, V., Kehrer, N., Vacchini, A., Chancellor, A., Contassot, E., Beshirova, A., Prota, G., Navarini, A., Mori, L. & De Libero, G. MR1 Gene and Protein Expression Are Enhanced by Inhibition of the Extracellular Signal-Regulated Kinase ERK. Cancer Immunology Research OF1–OF16 (2024) doi:10.1158/2326-6066.CIR-24-0110.

47. Narayan, V., Pion, E., Landré, V., Müller, P. & Ball, K. L. Docking-dependent ubiquitination of the interferon regulatory factor-1 tumor suppressor protein by the ubiquitin ligase CHIP. J Biol Chem 286, 607–619 (2011).

48. Rosain, J., Neehus, A.-L., Manry, J., Yang, R., Le Pen, J., Daher, W., Liu, Z., Chan, Y.-H., Tahuil, N., Türel, Ö., Bourgey, M., Ogishi, M., Doisne, J.-M., Izquierdo, H. M., Shirasaki, T., Le Voyer, T., Guérin, A., Bastard, P., Moncada-Vélez, M., Han, J. E., Khan, T., Rapaport, F., Hong, S.-H., Cheung, A., Haake, K., Mindt, B. C., Pérez, L., Philippot, Q., Lee, D., Zhang, P., Rinchai, D., Al Ali, F., Ahmad Ata, M. M., Rahman, M., Peel, J. N., Heissel, S., Molina, H., Kendir-Demirkol, Y., Bailey, R., Zhao, S., Bohlen, J., Mancini, M., Seeleuthner, Y., Roelens, M., Lorenzo, L., Soudée, C., Paz, M. E. J., González, M. L., Jeljeli, M., Soulier, J., Romana, S., L’Honneur, A.-S., Materna, M., Martínez-Barricarte, R., Pochon, M., Oleaga-Quintas, C., Michev, A., Migaud, M., Lévy, R., Alyanakian, M.-A., Rozenberg, F., Croft, C. A., Vogt, G., Emile, J.-F., Kremer, L., Ma, C. S., Fritz, J. H., Lemon, S. M., Spaan, A. N., Manel, N., Abel, L., MacDonald, M. R., Boisson-Dupuis, S., Marr, N., Tangye, S. G., Di Santo, J. P., Zhang, Q., Zhang, S.-Y., Rice, C. M., Béziat, V., Lachmann, N., Langlais, D., Casanova, J.-L., Gros, P. & Bustamante, J. Human IRF1 governs macrophagic IFN-γ immunity to mycobacteria. Cell 186, 621–645.e33 (2023).

49. Kim, S.-J. & Karamooz, E. MR1- and HLA-E-Dependent Antigen Presentation of Mycobacterium tuberculosis. International Journal of Molecular Sciences 23, 14412 (2022).

50. Cross, D. L., Layton, E. D., Yu, K. K. Q., Smith, M. T., Aguilar, M. S., Li, S., Wilcox, E. C., Chapuis, A. G., Mayanja-Kizza, H., Stein, C. M., Boom, W. H., Hawn, T. R., Bradley, P., Newell, E. W. & Seshadri, C. MR1-restricted T cell clonotypes are associated with “resistance” to *Mycobacterium tuberculosis* infection. JCI Insight 9, (2024).

51. Yang, R., Mele, F., Worley, L., Langlais, D., Rosain, J., Benhsaien, I., Elarabi, H., Croft, C. A., Doisne, J.-M., Zhang, P., Weisshaar, M., Jarrossay, D., Latorre, D., Shen, Y., Han, J., Ogishi, M., Gruber, C., Markle, J., Ali, F. A., Rahman, M., Khan, T., Seeleuthner, Y., Kerner, G., Husquin, L. T., Maclsaac, J. L., Jeljeli, M., Errami, A., Ailal, F., Kobor, M. S., Oleaga-Quintas, C., Roynard, M., Bourgey, M., Baghdadi, J. E., Boisson-Dupuis, S., Puel, A., Batteux, F., Rozenberg, F., Marr, N., Pan-Hammarström, Q., Bogunovic, D., Quintana-Murci, L., Carroll, T., Ma, C. S., Abel, L., Bousfiha, A., Santo, J. P. D., Glimcher, L. H., Gros, P., Tangye, S. G., Sallusto, F., Bustamante, J. & Casanova, J.-L. Human T-bet governs innate and innate-like adaptive IFN-γ immunity against mycobacteria. Cell 183, 1826 (2020).

52. Ussher, J. E., van Wilgenburg, B., Hannaway, R. F., Ruustal, K., Phalora, P., Kurioka, A., Hansen, T. H., Willberg, C. B., Phillips, R. E. & Klenerman, P. TLR signaling in human antigen-presenting cells regulates MR1-dependent activation of MAIT cells. Eur J Immunol 46, 1600–1614 (2016).

53. Lamichhane, R., Galvin, H., Hannaway, R. F., Harpe, S. M. de la, Munro, F., Tyndall, J. D., Vernall, A. J., McCall, J. L., Husain, M. & Ussher, J. E. Type I interferons are important co-stimulatory signals during T cell receptor mediated human MAIT cell activation. European Journal of Immunology 50, 178–191 (2020).

54. Pavlovic, M., Gross, C., Chili, C., Secher, T. & Treiner, E. MAIT Cells Display a Specific Response to Type 1 IFN Underlying the Adjuvant Effect of TLR7/8 Ligands. Front Immunol 11, 2097 (2020).

55. López-Rodríguez, J. C., Hancock, S. J., Li, K., Crotta, S., Barrington, C., Suárez-Bonnet, A., Priestnall, S. L., Aubé, J., Wack, A., Klenerman, P., Bengoechea, J. A. & Barral, P. Type I interferons drive MAIT cell functions against bacterial pneumonia. Journal of Experimental Medicine 220, e20230037 (2023).

56. Lazear, H. M., Schoggins, J. W. & Diamond, M. S. Shared and Distinct Functions of Type I and Type III Interferons. Immunity 50, 907–923 (2019).

57. Forero, A., Ozarkar, S., Li, H., Lee, C. H., Hemann, E. A., Nadjsombati, M. S., Hendricks, M. R., So, L., Green, R., Roy, C. N., Sarkar, S. N., von Moltke, J., Anderson, S. K., Gale, M. & Savan, R. Differential activation of the transcription factor IRF1 underlies the distinct immune responses elicited by type I and type III interferons. Immunity 51, 451–464.e6 (2019).

58. Goel, R. R., Kotenko, S. V. & Kaplan, M. J. Interferon lambda in inflammation and autoimmune rheumatic diseases. Nat Rev Rheumatol 17, 349–362 (2021).

59. Lavi, E., Suzumura, A., Murasko, D. M., Murray, E. M., Silberger, D. H. & Weiss, S. R. Tumor necrosis factor induces expression of MHC class I antigens on mouse astrocytes. Journal of Neuroimmunology 18, 245 (2002).

60. Venkatesh, D., Ernandez, T., Rosetti, F., Batal, I., Cullere, X., Luscinskas, F. W., Zhang, Y., Stavrakis, G., García-Cardeña, G., Horwitz, B. H. & Mayadas, T. N. Endothelial TNF receptor 2 induces IRF1 transcription factor-dependent interferon-β autocrine signaling to promote monocyte recruitment. Immunity 38, 1025–1037 (2013).

61. Tulli, L., Cattaneo, F., Vinot, J., Baldari, C. T. & D’Oro, U. Src Family Kinases Regulate Interferon Regulatory Factor 1 K63 Ubiquitination following Activation by TLR7/8 Vaccine Adjuvant in Human Monocytes and B Cells. Front. Immunol. 9, (2018).

62. Liu, A., Gong, P., Hyun, S. W., Wang, K. Z. Q., Cates, E. A., Perkins, D., Bannerman, D. D., Puché, A. C., Toshchakov, V. Y., Fang, S., Auron, P. E., Vogel, S. N. & Goldblum, S. E. TRAF6 Protein Couples Toll-like Receptor 4 Signaling to Src Family Kinase Activation and Opening of Paracellular Pathway in Human Lung Microvascular Endothelia*. Journal of Biological Chemistry 287, 16132–16145 (2012).

63. Harikumar, K. B., Yester, J. W., Surace, M. J., Oyeniran, C., Price, M. M., Huang, W.-C., Hait, N. C., Allegood, J. C., Yamada, A., Kong, X., Lazear, H. M., Bhardwaj, R., Takabe, K., Diamond, M. S., Luo, C., Milstien, S., Spiegel, S. & Kordula, T. K63-linked polyubiquitination of transcription factor IRF1 is essential for IL-1-induced production of chemokines CXCL10 and CCL5. Nat Immunol 15, 231–238 (2014).

64. Hoshino, K., Sasaki, I., Sugiyama, T., Yano, T., Yamazaki, C., Yasui, T., Kikutani, H. & Kaisho, T. Critical role of IkappaB Kinase alpha in TLR7/9-induced type I IFN production by conventional dendritic cells. J Immunol 184, 3341–3345 (2010).

65. Sánchez-Tarjuelo, R., Cortegano, I., Manosalva, J., Rodríguez, M., Ruíz, C., Alía, M., Prado, M. C., Cano, E. M., Ferrándiz, M. J., Campa, A. G. de la, Gaspar, M. L. & Andrés, B. de. The TLR4-MyD88 Signaling Axis Regulates Lung Monocyte Differentiation Pathways in Response to Streptococcus pneumoniae. Frontiers in Immunology 11, 2120 (2020).

66. Ioannidis, M., Cerundolo, V. & Salio, M. The Immune Modulating Properties of Mucosal-Associated Invariant T Cells. Front. Immunol. 0, (2020).

67. Davey, M. S., Morgan, M. P., Liuzzi, A. R., Tyler, C. J., Khan, M. W. A., Szakmany, T., Hall, J. E., Moser, B. & Eberl, M. Microbe-Specific Unconventional T Cells Induce Human Neutrophil Differentiation into Antigen Cross-Presenting Cells. The Journal of Immunology 193, 3704–3716 (2014).

68. Siebeler, R., de Winther, M. P. J. & Hoeksema, M. A. The regulatory landscape of macrophage interferon signaling in inflammation. Journal of Allergy and Clinical Immunology 152, 326–337 (2023).

69. Frucht, D. M., Fukao, T., Bogdan, C., Schindler, H., O’Shea, J. J. & Koyasu, S. IFN-γ production by antigen-presenting cells: mechanisms emerge. Trends in Immunology 22, 556–560 (2001).

70. Fenimore, J. & Young, H. A. Regulation of IFN-γ Expression. in Regulation of Cytokine Gene Expression in Immunity and Diseases (ed. Ma, X.) 1–19 (Springer Netherlands, 2016). doi:10.1007/978-94-024-0921-5_1.

71. Gomez, J. C., Yamada, M., Martin, J. R., Dang, H., Brickey, W. J., Bergmeier, W., Dinauer, M. C. & Doerschuk, C. M. Mechanisms of Interferon-γ Production by Neutrophils and Its Function during *Streptococcus pneumoniae* Pneumonia. Am J Respir Cell Mol Biol 52, 349–364 (2015).

72. Briard, B., Karki, R., Malireddi, R. K. S., Bhattacharya, A., Place, D. E., Mavuluri, J., Peters, J. L., Vogel, P., Yamamoto, M. & Kanneganti, T.-D. Fungal ligands released by innate immune effectors promote inflammasome activation during Aspergillus fumigatus infection. Nat Microbiol 4, 316–327 (2019).

73. Hinks, T. S. Mucosal-associated invariant T cells in autoimmunity, immune-mediated diseases and airways disease. Immunology 148, 1–12 (2016).

74. Pincikova, T., Parrot, T., Hjelte, L., Högman, M., Lisspers, K., Ställberg, B., Janson, C., Malinovschi, A. & Sandberg, J. K. MAIT cell counts are associated with the risk of hospitalization in COPD. Respiratory Research 23, 127 (2022).

75. Mock, J. R., Tune, M. K., Dial, C. F., Torres-Castillo, J., Hagan, R. S. & Doerschuk, C. M. Effects of IFN-γ on immune cell kinetics during the resolution of acute lung injury. Physiol Rep 8, e14368 (2020).

76. Ramana, C. V., DeBerge, M. P., Kumar, A., Alia, C. S., Durbin, J. E. & Enelow, R. I. Inflammatory impact of IFN-γ in CD8+ T cell-mediated lung injury is mediated by both Stat1-dependent and -independent pathways. Am J Physiol Lung Cell Mol Physiol 308, L650–L657 (2015).

77. Southworth, T., Metryka, A., Lea, S., Farrow, S., Plumb, J. & Singh, D. IFN-γ synergistically enhances LPS signalling in alveolar macrophages from COPD patients and controls by corticosteroid-resistant STAT1 activation. Br J Pharmacol 166, 2070–2083 (2012).

78. Williams, M., Todd, I. & Fairclough, L. C. The role of CD8 + T lymphocytes in chronic obstructive pulmonary disease: a systematic review. Inflamm. Res. 70, 11–18 (2021).

79. Zhu, X., Gadgil, A. S., Givelber, R., George, M. P., Stoner, M. W., Sciurba, F. C. & Duncan, S. R. Peripheral T Cell Functions Correlate with the Severity of Chronic Obstructive Pulmonary Disease. The Journal of Immunology 182, 3270–3277 (2009).

80. Briend, E., Ferguson, G. J., Mori, M., Damera, G., Stephenson, K., Karp, N. A., Sethi, S., Ward, C. K., Sleeman, M. A., Erjefält, J. S. & Finch, D. K. IL-18 associated with lung lymphoid aggregates drives IFNγ production in severe COPD. Respiratory Research 18, 159 (2017).

81. Seifert, L. L., Si, C., Saha, D., Sadic, M., de Vries, M., Ballentine, S., Briley, A., Wang, G., Valero-Jimenez, A. M., Mohamed, A., Schaefer, U., Moulton, H. M., García-Sastre, A., Tripathi, S., Rosenberg, B. R. & Dittmann, M. The ETS transcription factor ELF1 regulates a broadly antiviral program distinct from the type I interferon response. PLoS Pathog 15, e1007634 (2019).

82. Yang, L. & Ding, J. L. MEK1/2 Inhibitors Unlock the Constrained Interferon Response in Macrophages Through IRF1 Signaling. Frontiers in Immunology 10, 2020 (2019).

83. Najjar, I. & Fagard, R. STAT1 and pathogens, not a friendly relationship. Biochimie 92, 425–444 (2010).

84. Miorin, L. & Sanchez-Aparicio, M. T. SLiMs go viral! One more weapon against interferon. Cell Host & Microbe 30, 286–288 (2022).

85. Li, J., Yu, L., Shen, Z., Li, Y., Chen, B., Wei, W., Chen, X., Wang, Q., Tong, F., Lou, H., Chu, M. & Wei, L. miR-34a and its novel target, NLRC5, are associated with HPV16 persistence. Infection, Genetics and Evolution 44, 293–299 (2016).

86. Remoli, A. L., Marsili, G., Perrotti, E., Acchioni, C., Sgarbanti, M., Borsetti, A., Hiscott, J. & Battistini, A. HIV-1 Tat Recruits HDM2 E3 Ligase To Target IRF-1 for Ubiquitination and Proteasomal Degradation. mBio 7, 10.1128/mbio.01528-16 (2016).

87. Lewinsohn, D. M., Briden, A. L., Reed, S. G., Grabstein, K. H. & Alderson, M. R. *Mycobacterium tuberculosis*-reactive CD8+ T lymphocytes: the relative contribution of classical versus nonclassical HLA restriction. J. Immunol. 165, 925–30 (2000).

88. Wong, J., Korcheva, V., Jacoby, D. B. & Magun, B. E. Proinflammatory responses of human airway cells to ricin involve stress-activated protein kinases and NF-κB. American Journal of Physiology-Lung Cellular and Molecular Physiology 293, L1385–L1394 (2007).

89. Kulicke, C. A., Swarbrick, G. M., Ladd, N. A., Cansler, M., Null, M., Worley, A., Lemon, C., Ahmed, T., Bennett, J., Lust, T. N., Heisler, C. M., Huber, M. E., Krawic, J. R., Ankley, L. M., McBride, S. K., Tafesse, F. G., Olive, A. J., Hildebrand, W. H., Lewinsohn, D. A., Adams, E. J., Lewinsohn, D. M. & Harriff, M. J. Delivery of loaded MR1 monomer results in efficient ligand exchange to host MR1 and subsequent MR1T cell activation. Commun Biol 7, 1–13 (2024).

90. Hartmann, N., McMurtrey, C., Sorensen, M. L., Huber, M. E., Kurapova, R., Coleman, F. T., Mizgerd, J. P., Hildebrand, W., Kronenberg, M., Lewinsohn, D. M. & Harriff, M. J. Riboflavin Metabolism Variation among Clinical Isolates of *Streptococcus pneumoniae* Results in Differential Activation of Mucosal-associated Invariant T Cells. Am. J. Respir. Cell Mol. Biol. 58, 767–776 (2018).

91. Brinkman, E. K., Chen, T., Amendola, M. & van Steensel, B. Easy quantitative assessment of genome editing by sequence trace decomposition. Nucleic Acids Research 42, e168 (2014).

92. Conant, D., Hsiau, T., Rossi, N., Oki, J., Maures, T., Waite, K., Yang, J., Joshi, S., Kelso, R., Holden, K., Enzmann, B. L. & Stoner, R. Inference of CRISPR Edits from Sanger Trace Data. The CRISPR Journal 5, 123–130 (2022).

93. Li, K., Vorkas, C. K., Chaudhry, A., Bell, D. L., Willis, R. A., Rudensky, A., Altman, J. D., Glickman, M. S. & Aubé, J. Synthesis, stabilization, and characterization of the MR1 ligand precursor 5-amino-6-D-ribitylaminouracil (5-A-RU). PLOS ONE 13, e0191837 (2018).

94. Livak, K. J. & Schmittgen, T. D. Analysis of relative gene expression data using real-time quantitative PCR and the 2(-Delta Delta C(T)) Method. Methods 25, 402–8 (2001).

95. Heinzel, A. S., Grotzke, J. E., Lines, R. A., Lewinsohn, D. A., McNabb, A. L., Streblow, D. N., Braud, V. M., Grieser, H. J., Belisle, J. T. & Lewinsohn, D. M. HLA-E-dependent presentation of Mtb-derived antigen to human CD8+ T cells. J. Exp. Med. 196, 1473–81 (2002).

96. Muller, R. Y., Hammond, M. C., Rio, D. C. & Lee, Y. J. An Efficient Method for Electroporation of Small Interfering RNAs into ENCODE Project Tier 1 GM12878 and K562 Cell Lines. J Biomol Tech 26, 142–149 (2015).

97. Dreos, R., Ambrosini, G., Groux, R., Cavin Périer, R. & Bucher, P. The eukaryotic promoter database in its 30th year: focus on non-vertebrate organisms. Nucleic Acids Research 45, D51–D55 (2017).

98. Wasserman, W. W. & Sandelin, A. Applied bioinformatics for the identification of regulatory elements. Nat Rev Genet 5, 276–287 (2004).

99. Ambrosini, G., Praz, V., Jagannathan, V. & Bucher, P. Signal search analysis server. Nucleic Acids Res 31, 3618–3620 (2003).

