## Supplementary figures and images for "IFNγ regulates MR1 transcription and antigen presentation"

### Supplemental Figure 1

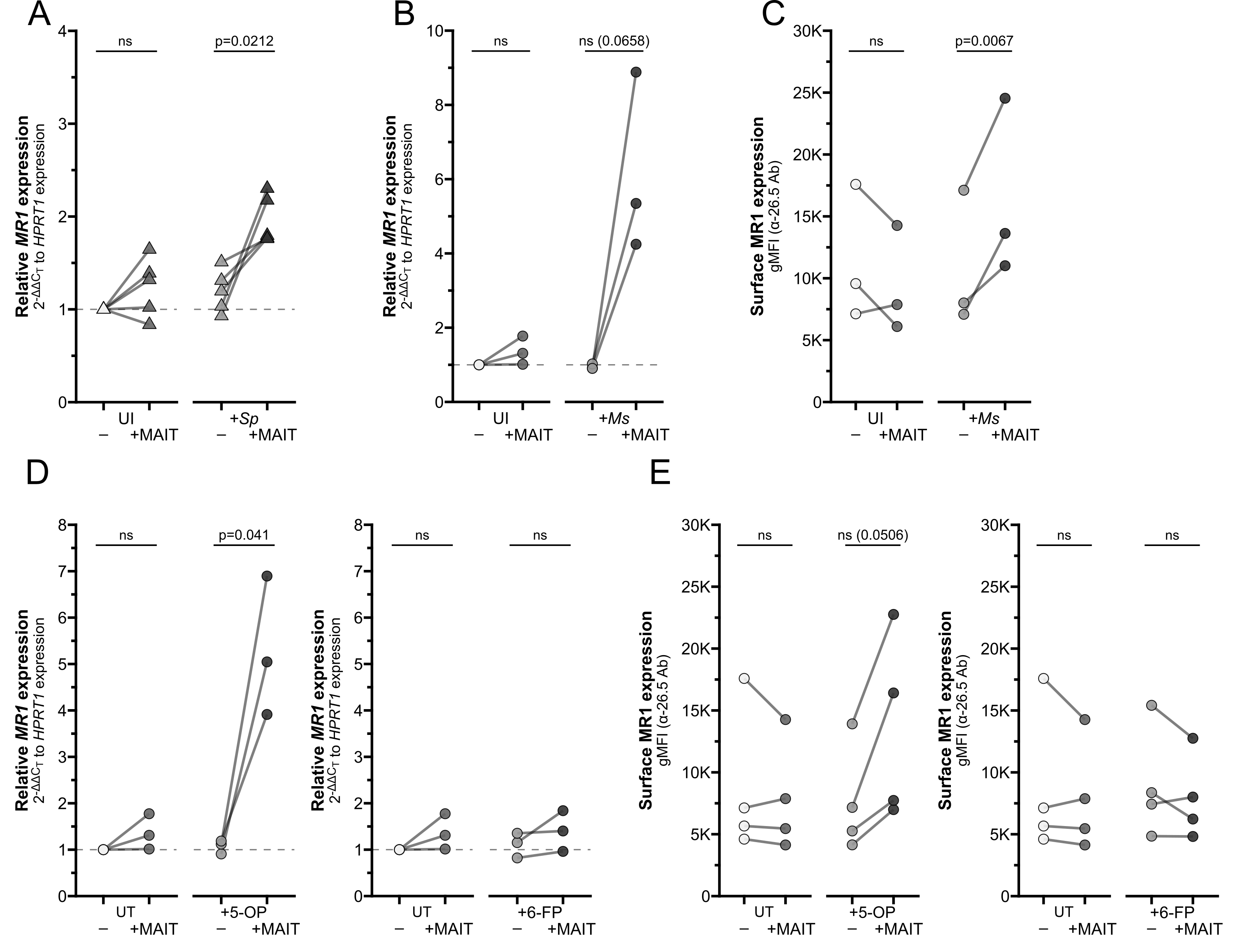

### Supplemental Figure 2

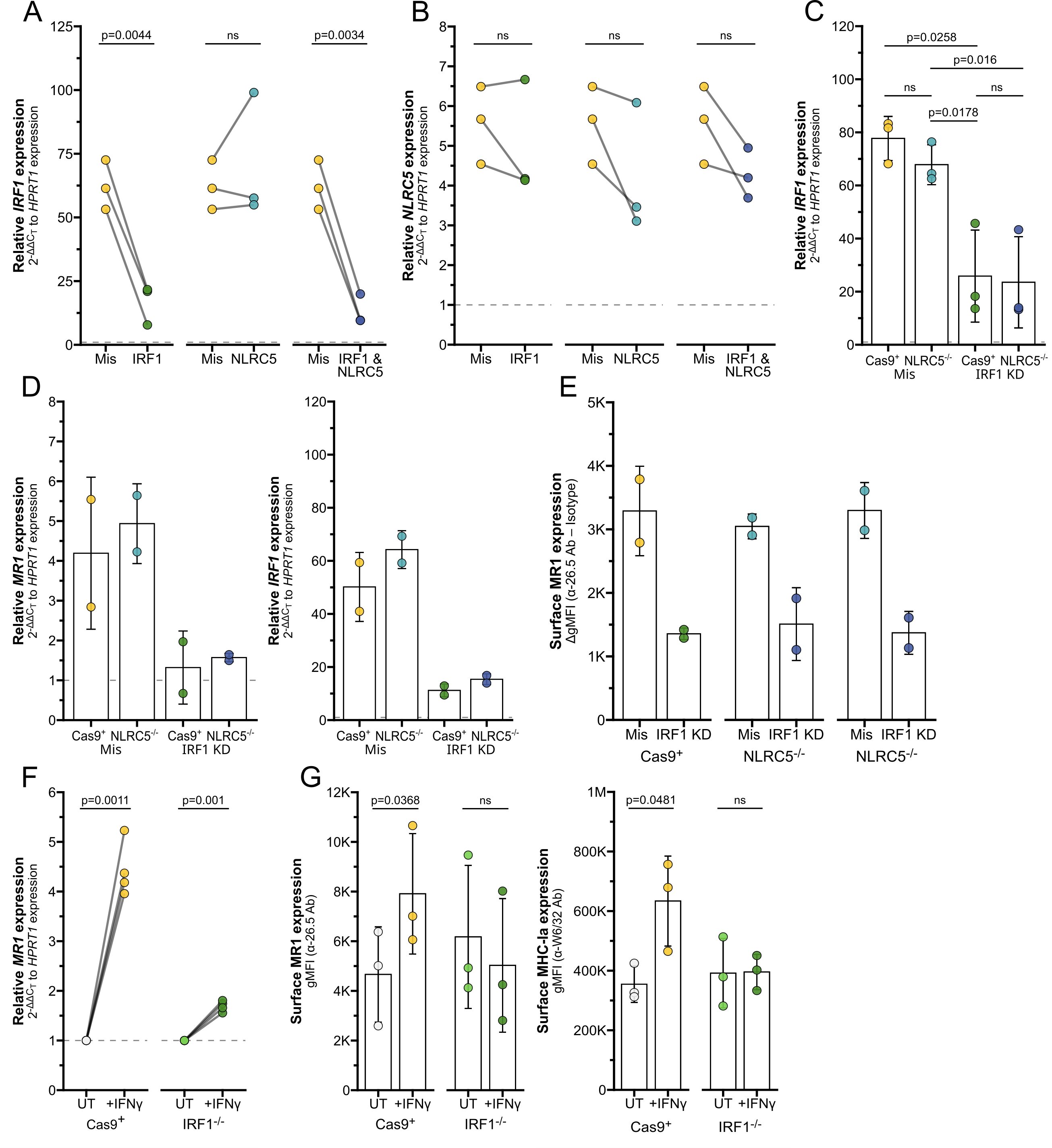

### Supplemental Figure 3

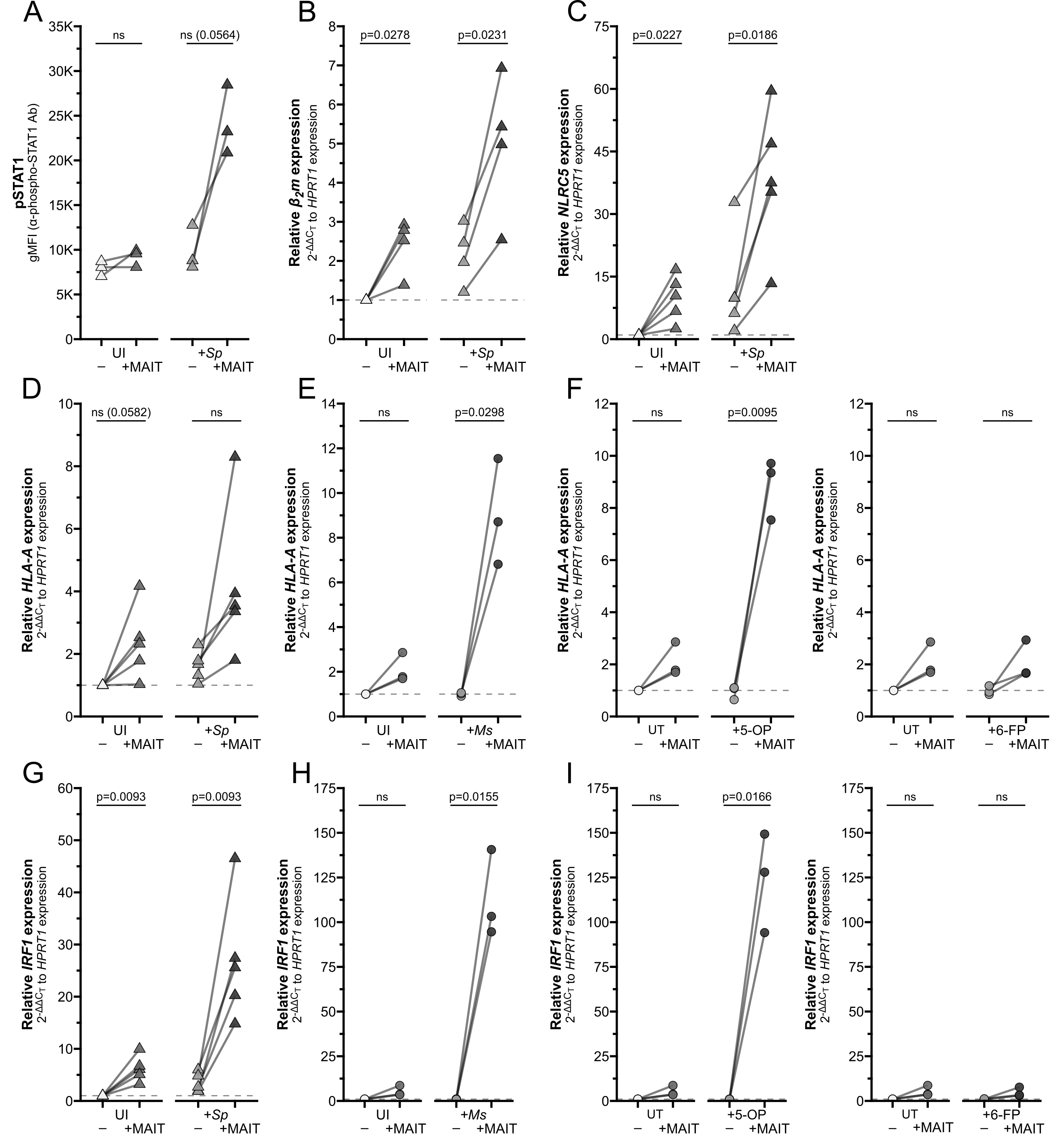

### Supplemental Figure 4

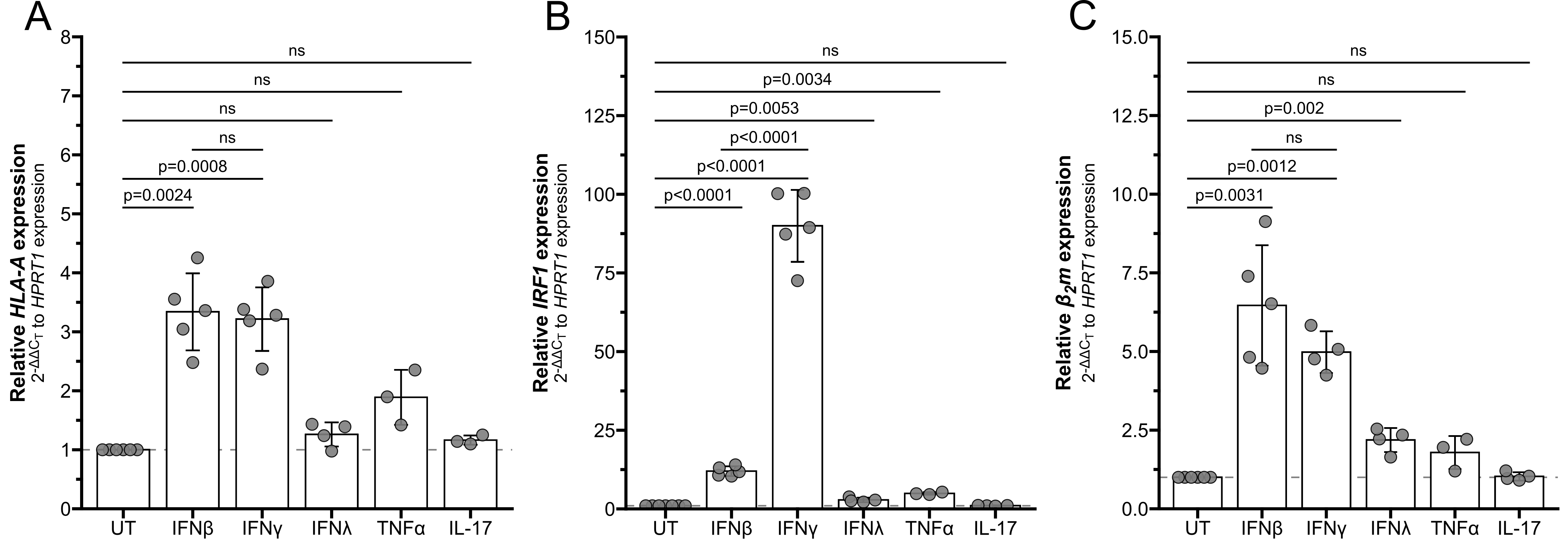
